# Conduit integrity is compromised during acute lymph node expansion

**DOI:** 10.1101/527481

**Authors:** Victor G. Martinez, Valeriya Pankova, Lukas Krasny, Tanya Singh, Ian J. White, Agnesska C. Benjamin, Simone Dertschnig, Harry L. Horsnell, Janos Kriston-Vizi, Jemima J. Burden, Paul H. Huang, Christopher J. Tape, Sophie E. Acton

**Affiliations:** Stromal Immunology Group, MRC Laboratory for Molecular Cell Biology, University College London, Gower Street, London, WC1E 6BT, United Kingdom; Division of Molecular Pathology, Institute of Cancer Research; 237 Fulham Road London, SW3 6JB, UK; Bioinformatics Image Core, MRC Laboratory for Molecular Cell Biology, University College London; London, WC1E 6BT, UK; Electron microscopy facility, MRC Laboratory for Molecular Cell Biology, University College London, London, WC1E 6BT, UK; UCL Institute of Immunity and Transplantation, University College London, London, NW3 2PF, UK; Cell Communication Lab, Department of Oncology, University College London Cancer Institute, 72 Huntley Street, London, WC1E 6DD, UK

**Keywords:** Dendritic cell, Fibroblastic reticular network, Conduit, Extracellular matrix, Lymph Node, Podoplanin, CLEC-2, LL5*β*

## Abstract

Lymph nodes (LNs) work as filtering organs, constantly sampling peripheral cues. This is facilitated by the conduit network, a parenchymal tubular-like structure formed of bundles of aligned extracellular matrix (ECM) fibrils ensheathed by fibroblastic reticular cells (FRCs). LNs undergo 5-fold expansion with every adaptive immune response and yet these ECM-rich structures are not permanently damaged. Whether conduit integrity and filtering functions are affected during cycles of LN expansion and resolution is not known. Here we show that the conduit structure is disrupted during acute LN expansion but FRC-FRC contacts remain intact. In homeostasis, polarised FRCs adhere to the underlying substrate to deposit ECM basolaterally. ECM production by FRCs is regulated by the C-type lectin CLEC-2, expressed by dendritic cells (DCs), at transcriptional and secretory levels. Inflamed LNs maintain conduit size-exclusion, but flow becomes leaky, which allows soluble antigens to reach more antigen-presenting cells. We show how dynamic communication between peripheral tissues and LNs changes during immune responses, and describe a mechanism that enables LNs to prevent inflammation-induced fibrosis.

**Highlights:**

- FRCs use polarized microtubule networks to guide matrix deposition
- CLEC-2/PDPN controls matrix production at transcriptional and post-transcriptional levels
- FRCs halt matrix production and decouple from conduits during acute LN expansion
- Conduits leak soluble antigen during acute LN expansion

## Introduction

Lymph node (LN) functional organisation is formed and maintained by the stromal cells. Once thought to only provide the necessary scaffolds to support the architecture of the tissue, LN stromal cells are now recognised as key players in immunity (1). Four main populations of stromal cells can be defined in LNs: podoplanin (PDPN)-CD31+ blood endothelial cells (BECs), PDPN+CD31+ lymphatic endothelial cells (LECs), PDPN+CD31-fibroblastic reticular cells (FRCs) and PDPN-CD31-double-negative cells (DN) (2). Among these, the FRC population is the most abundant stromal subset. A recent scRNAseq analysis showed that FRCs may contain different subpopulations with specific locations and functions (3). FRCs form a connected 3D network that spans the T-cell area and interfollicular regions of LNs. FRCs regulate lymphocyte homeostasis (4, 5) and induction of peripheral tolerance (6–8). Furthermore, contraction through the FRC network also regulates the immune response at a whole organ level by controlling LN size. Interactions between FRC and C-type lectin-like receptor 2 (CLEC-2)+ migratory dendritic cells (DCs) transiently inhibit PDPN-dependent actomyosin contractility during the acute phase of the immune response (9, 10), allowing rapid LN expansion.

LNs also function as filters for lymph-born antigens (11, 12). Soluble antigens reach the LN first in a wave of draining-type diffusion ahead of a secondary wave of migratory antigen-presenting cells. Collected by lymphatic capillaries in the peripheral tissue, the lymph converges in afferent lymphatic vessels that merge with the LN capsule, and flows within the subcapsular sinus (SCS). The lymph percolates through trabecular and cortical sinuses that flow into the LN medulla before leaving via efferent lymphatic vessels. Subcapsular and medullary sinuses are populated by macrophages that represent a first line of defence in LNs against pathogens (13). A sample of low molecular weight molecules (<70kDa) (14–17) is permitted to flow directly through the LN parenchyma within an intricate tubular system termed the conduit network (17), while the SCS and the associated macrophages prevent entry of viruses, bacteria, nanoparticles, and apoptotic cells. The conduit network is composed of highly bundled and aligned extracellular matrix (ECM) components enwrapped by FRCs (16, 18, 19). While we are starting to understand how the cellular part of the conduit, the FRC network, react during LN expansion, it is not yet known how the non-cellular ECM components are remodelled and how rapid expansion of the LN may affect the function of the conduit. In other contexts, tissue remodelling goes hand-in-hand with inflammation. Injury-induced loss of ECM is rapidly replenished by biogenesis and crosslinking (20). Chronic inflammation, as occurs in cancer or some viral infections, induces deregulation of this process leading to fibrosis (21).

LN fibrosis can occur in some cases of chronic viral-infection or tumour-draining LNs (22–25), however, more commonly, LNs undergo a virtually unlimited number of inflammatory episodes throughout an individual’s lifetime and LN fibrosis does not occur. In LNs, FRCs are the main producers of ECM (19). Therefore, we hypothesised that a specific mechanism must be in place in order to avoid the accumulation of aberrant ECM in lymphoid tissue.

In this study we focussed our investigations on ECM remodelling by FRCs during LN expansion, and the interconnection between the cellular and ECM components of the conduit network. We demonstrate a loss of ECM components of the conduit during acute LN expansion. We show that CLEC-2 binding to PDPN+ FRCs modulates ECM production at both mRNA and protein levels. Furthermore, the CLEC-2/PDPN axis regulates polarised microtubule organisation in FRCs to control ECM deposition. The resulting loss of conduit integrity alters intranodal flow of small molecules in inflamed LNs, allowing enhanced capture by LN resident myeloid cells.

## Results

### Extracellular matrix components of the conduit are reduced during acute LN expansion

To ask how ECM structures were maintained and remodelled during acute LN expansion, a period of rapid tissue growth, we first examined LN ECM structures in the steady state. Using the passive clarity technique (PACT) (26), we imaged collagen IV in intact naïve inguinal LNs (Figure 1A, Supplementary video 1). This abundant basement membrane protein surrounded the LN vasculature and formed an intricate 3D connected network spanning the whole LN parenchyma (16, 18, 19), corresponding to the conduit network. Electron microscopy revealed the detail of condensed fibrillar bundles consisting >200 collated fibres of ECM enwrapped by FRCs (Figure 1B). Co-staining of the basement membrane protein laminin and the FRC marker PDPN confirmed that in LNs parenchyma, ECM structures are associated with the FRC network (27), and vasculature (Figure 1C).

**Fig. 1.**
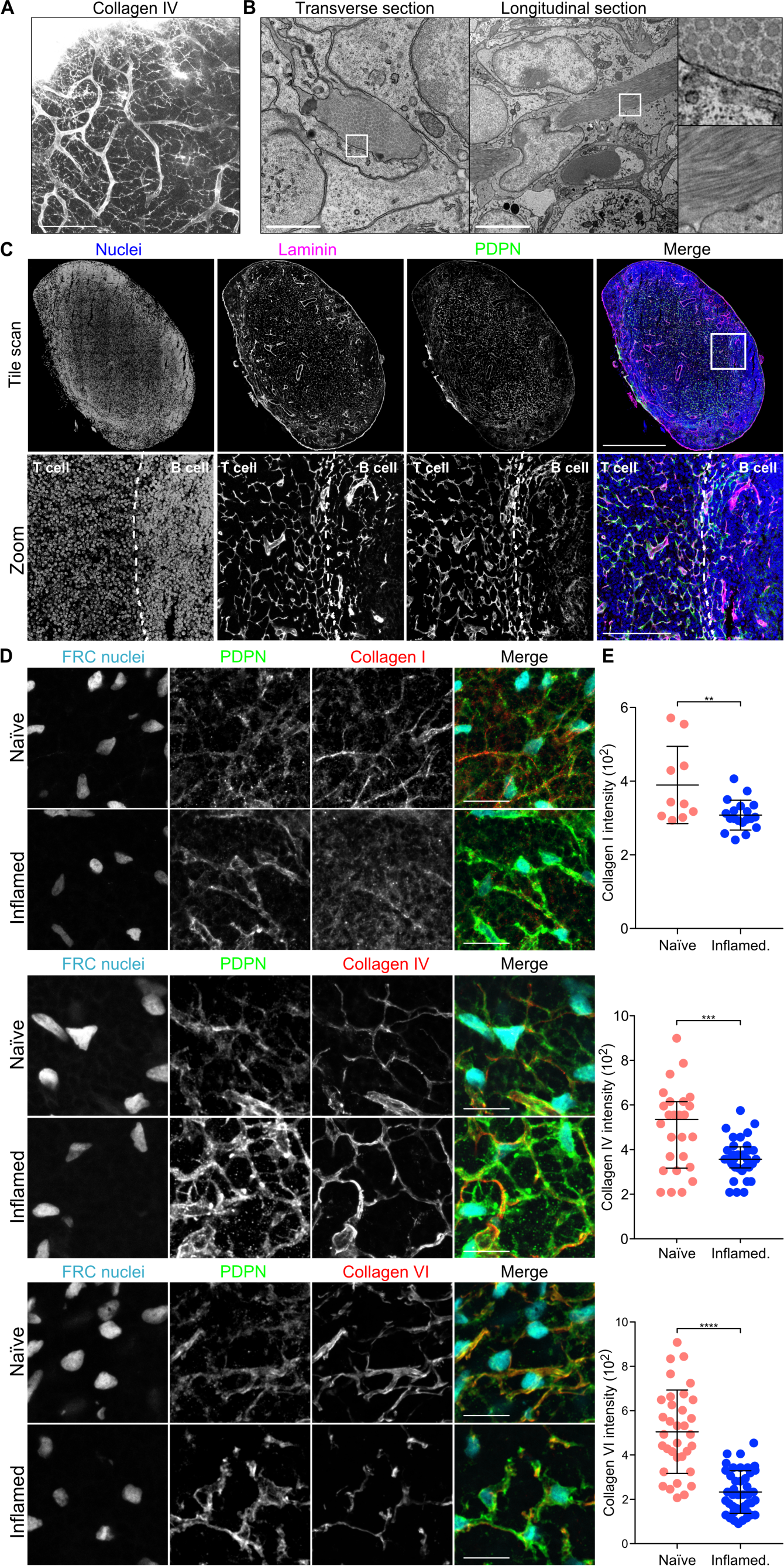
Conduit composition changes during LN expansion. A) Immunofluorescence of intact whole naïve lymph nodes stained for collagen IV using PACT. A maximum Z stack projection is shown. Scale bar represents 100 microns. B) Transmission electron microscopy of naïve LNs. Two sections and the indicated zoomed areas (right panels) are shown. The scale bars represent 5 microns. C) Immunoflu-orescence of 20 microns thick cryosection of a naïve LN. Maximum z stack projections of a tile scan (top) and zoomed area (bottom) are shown. Boundary between T and B cell areas is delineated by a discontinued line. The scale bars represent 500 microns (tile scan) and 100 microns (zoom). D-E) Immunofluorescence of 10 microns thick cryosection of naïve and inflamed LNs from PDGFR*α*H2B-GFP mice immunized with CFA/OVA. D) Maximum z stack projections of representative images are shown. The scale bars represent 20 microns. E) Quantification of the indicated ECM components within the PDPN network. Each dot represents the median grey intensity of a different region of interest (n=3). Error bars represent mean and SD. **P<0.005, ***P<0.0005, unpaired t test.

We immunised PDGFR*α*H2B-GFP mice with ovalbumin emulsified in complete Freund’s adjuvant (CFA/OVA), in which FRCs are identified with nuclear GFP (9), and compared to naïve LNs the structure and density of several collagen components after 5 days of LN expansion. Collagens I, IV and VI, (16, 18, 19), were abundant in naïve LNs and almost completely filled the FRC network. However, despite that the FRC network remained intact and connected, these structures were either less prominent or partially disrupted 5 days post immunisation (Figure 1D). All three collagens were reduced within the PDPN+ cellular network (Figure 1E), indicating that while the FRC cellular network remained connected and intact, the accompanying ECM forming the conduit remained associated with the FRCs but were no longer replete.

### CLEC-2 binding regulates ECM production by FRCs

FRCs rapidly change their morphology and network architecture in response to CLEC-2+ migratory DCs (9). We hypothesised that the remodelling of the cellular network may also affect the remodelling of the associated ECM downstream of the same DC/stromal contacts. We stimulated FRCs in vitro with CLEC-2-Fc recombinant protein and compared transcriptional profiles by RNAseq (Figure 2 and Figure S1). Bulk analysis of the transcriptomic data comparing 6hr and 24hr CLEC-2-Fc treatment revealed that CLEC-2-Fc induced a transient and largely reversible gene regulation response in FRCs (Figure S1A), similar transient kinetics to the inhibition of PDPN-dependent contractility (9). Gene ontology analysis (28, 29) showed that genes encoding proteins in the extracellular space/region were most enriched (Figure 2A). Indeed, using the matrisome database (30–32) of all ECM proteins and associated factors, we found that FRCs expressed 570/743 matrisome genes in vitro, of which 75 (13%) were regulated >2-fold 6 hr after CLEC-2-Fc binding (Figure 2B and Figure S1B).

**Fig. 2.**
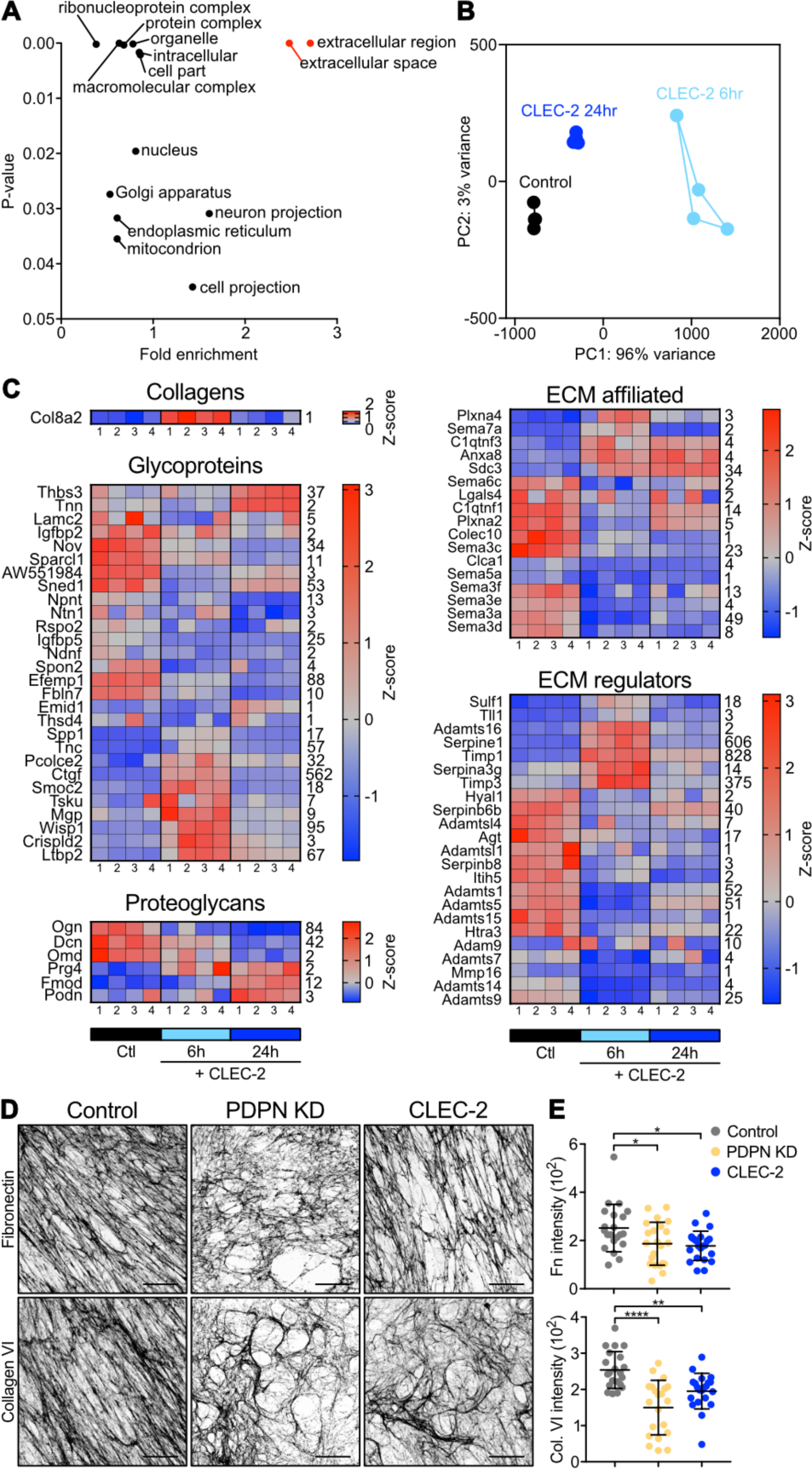
Effects of CLEC-2 on ECM production by FRCs. A-C) Gene expression by RNAseq in control FRCs treated with CLEC-2-Fc for 6 and 24 hours. A) Genes regulated by CLEC-2-Fc more or equal than 2-fold were subjected for statistical overrepresentation test of gene ontology analysis for cellular components. Each dot represents a cellular component significantly enriched, by binomial analysis. B) CLEC-2-Fc-regulated matrisomal gene cluster in a PCA space. C) Heatmap of matrisomal genes regulated by CLEC-2-Fc, clustered according to expression pattern. Four replicates for each condition is shown. Color code represents z-scores and the row average for each gene is indicated (right hand side of the heatmap). D) In vitro FRC cell line-derived matrices decellularized and stained for fibronectin (top) and collagen VI (bottom). Maximum z stack projections of representative images are shown. The scale bars represent 20 microns. E) Median grey intensity for the indicated ECM components. Each dot represents a different region of interest (n=3). Error bars represent mean and SD. *P<0.05**P<0.005, ***P<0.0005, one-way ANOVA, Tukey’s multiple comparisons test.

FRCs regulated 35 core matrisome genes (>2-fold) in response to CLEC-2-Fc, including one collagen (Col8a2), 23 glycoproteins, and 6 proteoglycans (Figure 2C). The down-regulated glycoproteins were mostly associated with cell-matrix adhesion and migration, including Nov, Sparcl1, Ntn1, Igfbp5, Ndnf, Spon2, Efemp1, Fbln7 (33–40). Glycoprotein genes induced had more pleiotropic roles, such us growth factor signalling (Ctgf, Tsku, Wisp1 and Ltbp2) (41–44) or immunomodulation (Spp1, Tnc, Crispld2) (45–47). Regulation of proteoglycan expression by CLEC-2-Fc was more evident at 24 hours, suggesting that these may be indirectly regulated; eg. CTGF/CCN2 represses Ogn, Dcn and Omd (48). CLEC-2-Fc increased expression of Prg4, which inhibits synoviocyte cell/matrix adhesion (49).

Most of the 17 ECM affiliated genes regulated by CLEC-2-Fc were linked to cytoskeleton regulation (50–52) (Figure 2C), including members of the semaphorin-plexin system which provide guidance cues for migration (52). Known to inhibit axonal growth (52), expression of Sema6c, Sema5a, Sema3f, Sema3e, Sema3a, Sema3d were reduced upon CLEC-2-Fc treatment, hinting that FRCs may spread using similar mechanisms. Of note, CLEC-2-Fc induced expression of Sema7a, which represses ECM production in other fibroblasts (53). CLEC-2-Fc regulation of 23 ECM regulators (Figure 2C) mainly affected protease inhibitors, including upregulation of Serpine1, Timp1 and Timp3, key in negative regulation of MMP activity (54, 55). Also upregulated, Sulf1 and Tll1 are involved in ECM biogenesis (56, 57). CLEC-2-Fc repressed the expression of several ECM regulator genes with prominent roles in ECM degradation: Hyal1 (Hyaluronidase-1) (58), Agt (SERPINA8/angiotensinogen) (59), Htra3 (60, 61), Adamts1 (62), Adamts5 (63), Adamts7 (64), Adamts9 (65), Adamts15 (66), Adam9 (67) and Mmp16 (68).

These data indicate that FRCs can substantially alter their transcription profile following CLEC-2 binding and that transcriptional regulation may play an important role in ECM remodelling and cell-matrix adhesion in FRCs. Furthermore, induction of protease inhibitors plus repression of proteases suggest that the observed loss of ECM within the conduit during LN expansion (Figure 1D) is unlikely to be due to degradation. Further, since we observed that collagens (I, IV and VI) are reduced in vivo in inflamed LNs (Figure 1D), but were not transcriptionally regulated by CLEC-2, this transcriptional regulation alone cannot fully explain the reduced ECM observed (Figure 1D).

To investigate whether the CLEC-2/PDPN signalling axis regulates ECM production at the protein level we undertook proteomic analysis on FRC-derived matrices in vitro (Figure S2). We generated CLEC-2-Fc-secreting FRCs to allow constant CLEC-2 stimulation and compared to PDPN-depleted FRCs (PDPN KD) (9) and a control FRC cell line. Mass spectrometry analysis detected a similar number of proteins in all three FRC cell lines of which 96 proteins were matrisomal proteins with a 90 % overlap among samples (Figure S2A). PDPN depletion phenocopies the same loss of contractility induced by CLEC-2 (9), in contrast, when comparing ECM protein production, PDPN KD FRCs appeared quantitatively very different from either control or CLEC-2-expressing FRCs. Among the proteins statistically changed (Figure S2B), PDPN KD FRC-derived matrices showed an overall reduction in ECM components, whereas CLEC-2-expressing FRCs and controls were more closely aligned (Figure S2C). This suggests that loss of PDPN expression is not equivalent to CLEC-2 modulation of PDPN function in the case of matrix production.

While the CLEC-2/PDPN signalling axis certainly influenced matrix transcription (Figure 2A-C) and protein production (Figure S2), how these changes translated to fibril formation, relevant to conduit remodelling in vivo, was still unclear. Staining of FRC-derived matrices for fibronectin and collagen VI showed that ECM structures formed by CLEC-2-expressing FRCs appeared disorganised compared to controls, with lower alignment and large empty spaces and lower median intensity of matrix fibres (Figure 2D and 2E). Interestingly, in this functional assay, PDPN KD FRCs phenocopied the effect of CLEC-2-Fc in matrix deposition and organisation. Together, these experiments demonstrate that CLEC-2/PDPN signalling regulates ECM remodelling at multiple levels, gene expression, protein production and fibril arrangement. These important in vitro results also show that PDPN expression by FRCs is a key requirement for FRCs to produce, deposit, and align ECM components, and that this process is disrupted by CLEC-2.

### Signalling cascades regulated by CLEC-2 in FRCs

The above results suggest that additional cellular mechanisms are likely to regulate ECM remodelling in FRCs. To address the CLEC-2/PDPN-dependent signalling cascades controlling ECM organisation, we performed unbiased phosphoproteomics analysis of FRCs by tandem mass tag mass spectrometry (TMT-MS) (69) (Figure 3A). Control FRCs were stimulated with CLEC-2-Fc for 15 min or 24 hr to capture immediate and late signalling responses. As expected, total protein levels did not differ significantly following treatment (Figure 3B). However, phosphoproteome analysis revealed that CLEC-2-Fc induced a rapid, and transient signalling response in FRCs (Figure 3C). At 15 min, 400 phosphorylation sites were regulated by CLEC-2-Fc (Figure 3D) corresponding to 77 proteins. In contrast, after 24 hr, only 7 phosphorylation sites corresponding to 6 proteins were regulated compared to controls, confirming the transient and reversible nature of responses to CLEC-2/PDPN engagement (Figure 3D).

**Fig. 3.**
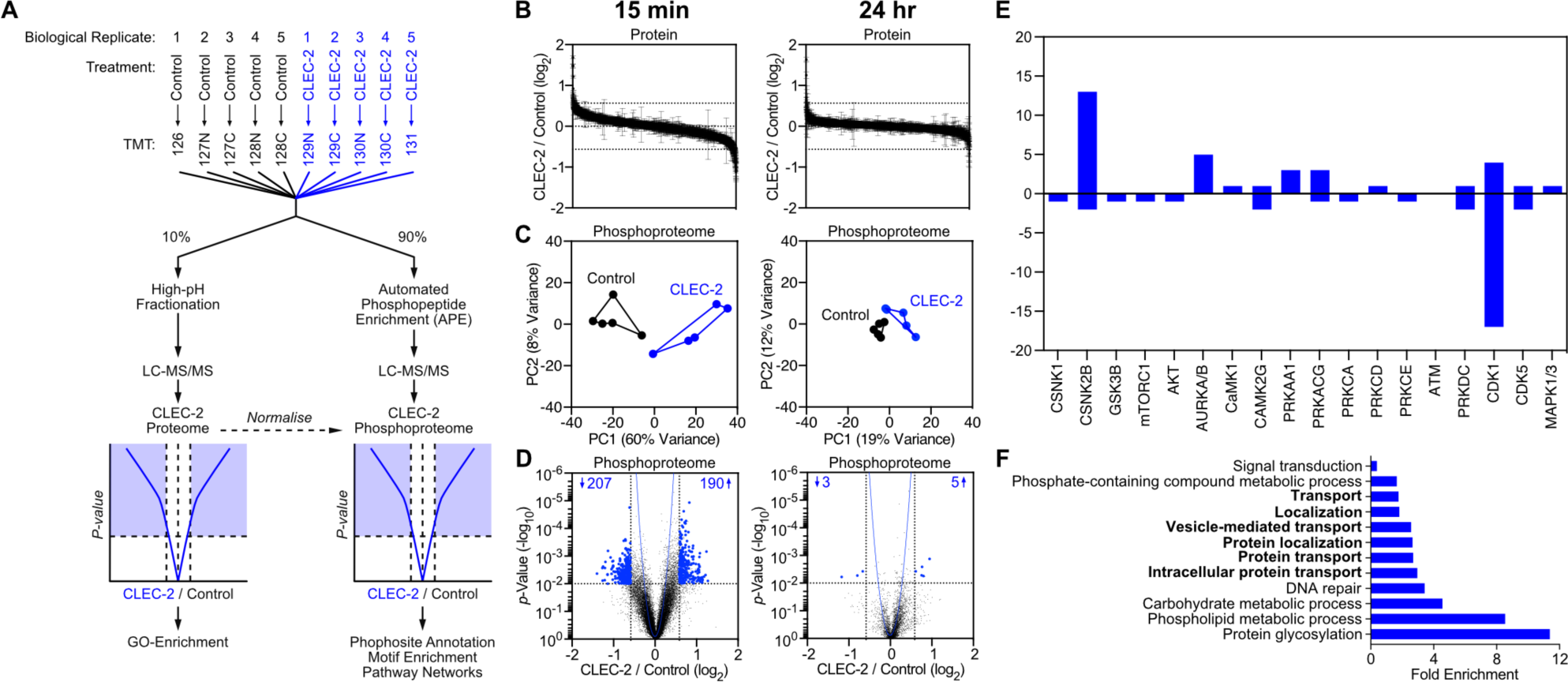
Phosphoproteomics of CLEC-2-treated FRCs. A) Control (untreated) and CLEC-2-Fc-treated FRCs cell lysates were isobarically labeled with tandem-mass tags (TMT) (126–131 mass-to-charge ratio [m/z]), mixed, and subjected to automatic phosphopeptide enrichment (APE) (n = 5). TMT-phosphopeptides were analyzed by high-resolution LC-MS/MS and normalized to total protein level changes. B) Waterfall graphs showing proteome regulation by CLEC-2-Fc. C) Control and CLEC-2-Fc-treated phosphoproteomes cluster in a PCA space. D) Volcano plots showing statistical regulation of the CLEC-2-Fc-treated FRC’s phosphoproteome (n = 5, two-tailed t test, Gaussian regression). Number of phosphosites considered statically different are indicated. E) Empirical parent kinase analysis. Bars represent number of targets for each putative parent kinases manually assigned to hits. Positive and negative values mean higher or lower phosphorylation in CLEC-2-Fc-treated cells. F) Statistical overrepresentation test of gene ontology analysis for biological processes. Each bar represents a biological process significantly enriched, by binomial analysis.

In order to elucidate signalling cascades regulated by CLEC-2, we performed kinase target analysis (15 min dataset). We found that CSNK2B and CDK1 regulated the highest number of predicted targets (Figure 3E). Gene ontology analysis of hits (28, 29) highlighted intracellular protein transport pathways (Figure 3F), of relevance to transport and deposition of cargo such as ECM components. However, using a GFP-based assay we found no reduction in protein secretion in either CLEC-2-Fc-treated or PDPN KD FRCs (Figure S3). Nevertheless, the impaired ECM deposition observed in both CLEC-2-Fc-treated and PDPN KD FRCs (Figure 2D) prompted a closer look at the phosphoproteomic data. Secretion of large cargo proteins like ECM components requires vesicle transport via cytoskeletal structures such as the microtubule network (70). We found that several key regulators of microtubule function were post-translationally modified by CLEC-2-Fc stimulation (Supplementary table 1), including Cytoplasmic linker protein 170 (CLIP-170), cytoplasmic dynein heavy chain 1, and pleckstrin homology-like domain family B member 2 (LL5*β*). While the direct function of these regulatory sites are not previously described, these data presented strong evidence that CLEC-2 altered the organisation of microtubules in FRCs, a possible regulatory mechanism of ECM deposition in LNs.

### CLEC-2 binding controls microtubule organization in FRCs via LL5*β*

LL5*β* forms complexes at the basal pole of epithelial cells attaching plus ends of microtubules to the cell cortex, forming a secretory pathway for local and polarised exocytosis (70–73). We examined the role of LL5*β* in ECM deposition by FRCs. We found that both CLEC-2-Fc treatment and PDPN KD reduced LL5*β* protein and mRNA levels in FRCs (Figure 4A and 4B). We attempted, unsuccessfully to overexpress a phosphomimetic mutant (LL5*β* S465E) in FRCs, leading us to hypothesise that phosphorylation of LL5*β* at S465 may target LL5*β* for degradation.

**Fig. 4.**
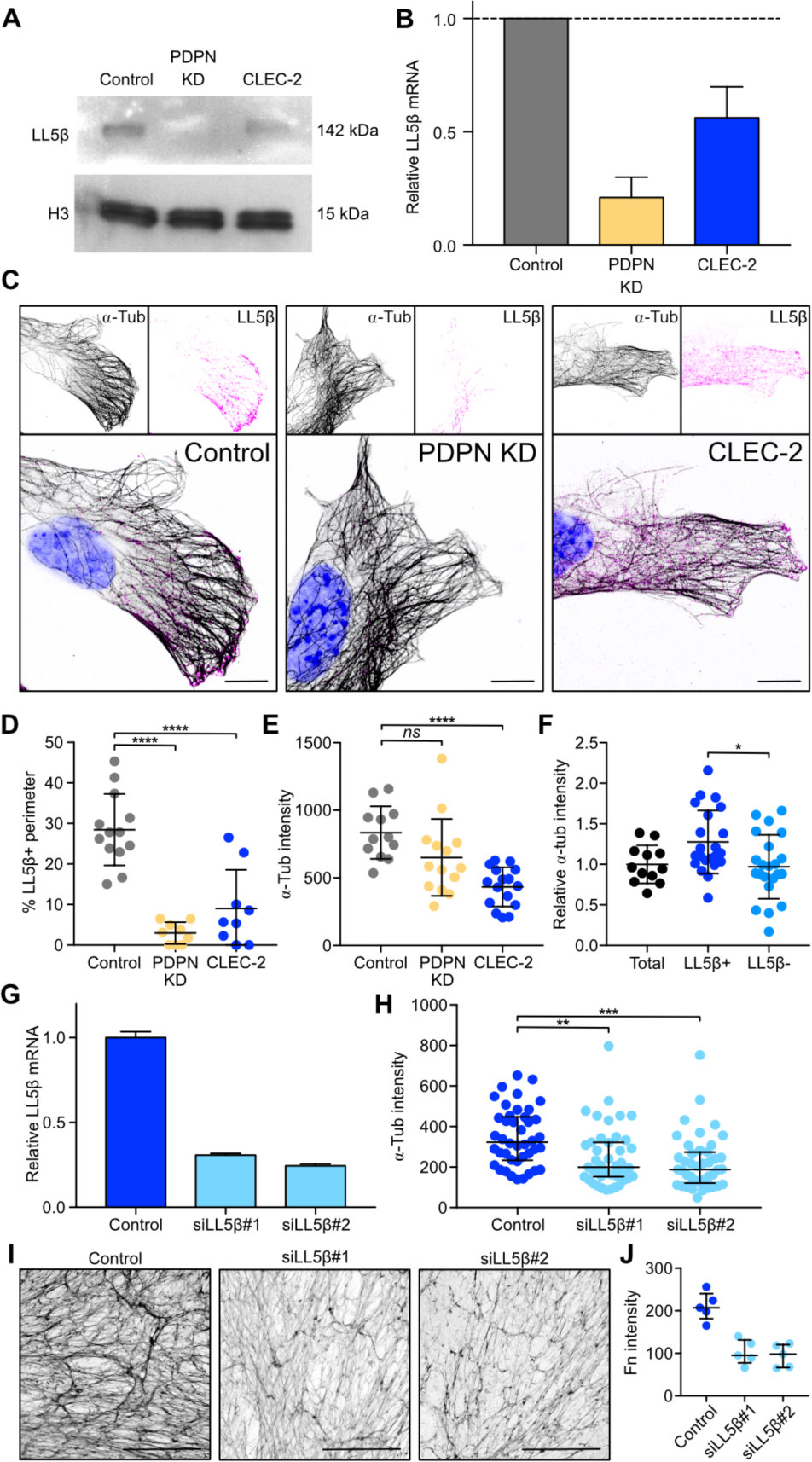
Regulation of microtubule organization by CLEC-2 via LL5*β*. A) Protein blots showing levels of LL5*β* in FRC cell lines. Histone 3 was used as a loading control. B) Expression of LL5*β* mRNA relative to control FRCs by qPCR. Error bars represent mean and SD (n=2). C) Immunofluorescence of FCR cell lines in culture. Maximum z stack projections of representative images are shown. The scale bars represent 10 microns. D) Quantification of LL5*β* coverage in FRC cell lines as a percentage of total perimeter. Each dot represents a cell (n=2). Error bars represent mean and SD. ****P<0.00005, one-way ANOVA, Tukey’s multiple comparisons test. E) *α*-tubulin intensity in the cortical area (10 microns deep) of FRC cell lines. Each dot represents a cell (n=2). Error bars represent mean and SD. ****P<0.00005, one-way ANOVA, Tukey’s multiple comparisons test. NS, not significant. F) *α*-tubulin intensity in the cortical area (10 microns deep) in LL5*β*-positive and -negative areas in control FRCs relative to total area. Black dots represent cells, blue dots represent cell areas (n=2). Error bars represent mean and SD. *P<0.05, one-way ANOVA, Tukey’s multiple comparisons test. G) Expression of LL5*β* mRNA by qPCR in LL5*β* siRNA transfected FRCs relative to control transfected FRCs (n=2). Error bars represent mean and SD. H) *α*-tubulin intensity in the cortical area (10 microns deep) of transfected FRCs. Each dot represents a cell (n=2). Error bars represent mean and SD. **P<0.005, ***P<0.0005, one-way ANOVA, Tukey’s multiple comparisons test. I) In vitro cell-derived matrices from transfected FRCs, decellularized and stained for fibronectin. Maximum z stack projections are shown. The scale bars represent 100 microns. J) Median grey intensity for fibronectin staining. Each dot represents a different region of interest. Error bars represent mean and SD.

Control FRCs clustered LL5*β* at the cell periphery (Figure 4C), however, this accumulation was absent in PDPN KD FRCs and more cytoplasmic in CLEC-2-Fc-treated cells (Figure 4C and 4D). The reduced cortical localisation of LL5*β* coincided with lower density of microtubules at the cell periphery in PDPN KD and CLEC-2-Fc treated cells (Figure 4E). We confirmed the colocalization of LL5*β* with cortical microtubules in control FRCs, where LL5*β*-positive areas presented higher microtubule density compared to areas lacking LL5*β* (Figure 4F). To ask if LL5*β* was required for microtubule attachment to the cortex in FRCs, we silenced LL5*β* expression using siRNA (Figure 4G) which resulted in a corresponding loss of microtubules from the periphery (Figure 4H). Importantly, LL5*β*-silenced FRCs also showed significantly reduced matrix deposition (Figure 4I and 4J), phenocopying the disrupted matrix generated following ei-ther CLEC-2-Fc treatment or PDPN depletion (Figure 2D), and confirming that LL5*β* is necessary for ECM deposition in FRCs.

### Loss of FRC adhesion and reorganisation of microtubule networks

LL5*β* is recruited to mature focal adhesion complexes, which require Rho-kinase (ROCK)/myosin-mediated contractility (73, 74). Since the CLEC-2/PDPN signalling axis inhibits actomyosin contractility in FRCs (9), we predicted it might also alter FRC adhesion to the underlying conduit and therefore inhibit the localisation of LL5*β* and microtubules to the cell cortex. We compared the structure and localisation of focal adhesions (p-paxillin) and LL5*β* between FRCs cell lines. CLEC-2-Fc-treated and PDPN KD cells presented significantly shorter focal adhesions (Figure 5A and 5B). This was phenocopied by the direct inhibition of ROCK (Y-27632) (Figure 5A and 5B). In controls, LL5*β* clustered directly adjacent to elongated mature focal adhesions (73) (Figure 5A and 5B). However, when focal adhesion maturation is disrupted, there is a concordant loss of LL5*β* clustering, linking actomyosin contractility, cell-matrix adhesion, and LL5*β* recruitment in an integrated mechanism (Figure 5A, 5B). To test these linked outcomes in a more physiological assay, we stimulated FRCs with either control or CD11cΔCLEC-2 bone marrow-derived dendritic cells (BMDCs). Cultured alone, FRCs displayed prominent f-actin stress fibres and mature elongated focal adhesions to which microtubules bundles docked in abundance (Figure 5C and 5D). Interaction with control (CLEC-2+) BMDCs induced loss of actin stress fibres, shorter focal adhesions and lower microtubule density at the periphery (Figure 5C and 5D). This change in FRC morphology and function was not observed with CD11cΔCLEC-2 BMDCs (Figure 5C and 5D), demonstrating that DC-induced inhibition of actomyosin contractility and microtubule localisation requires CLEC-2.

**Fig. 5.**
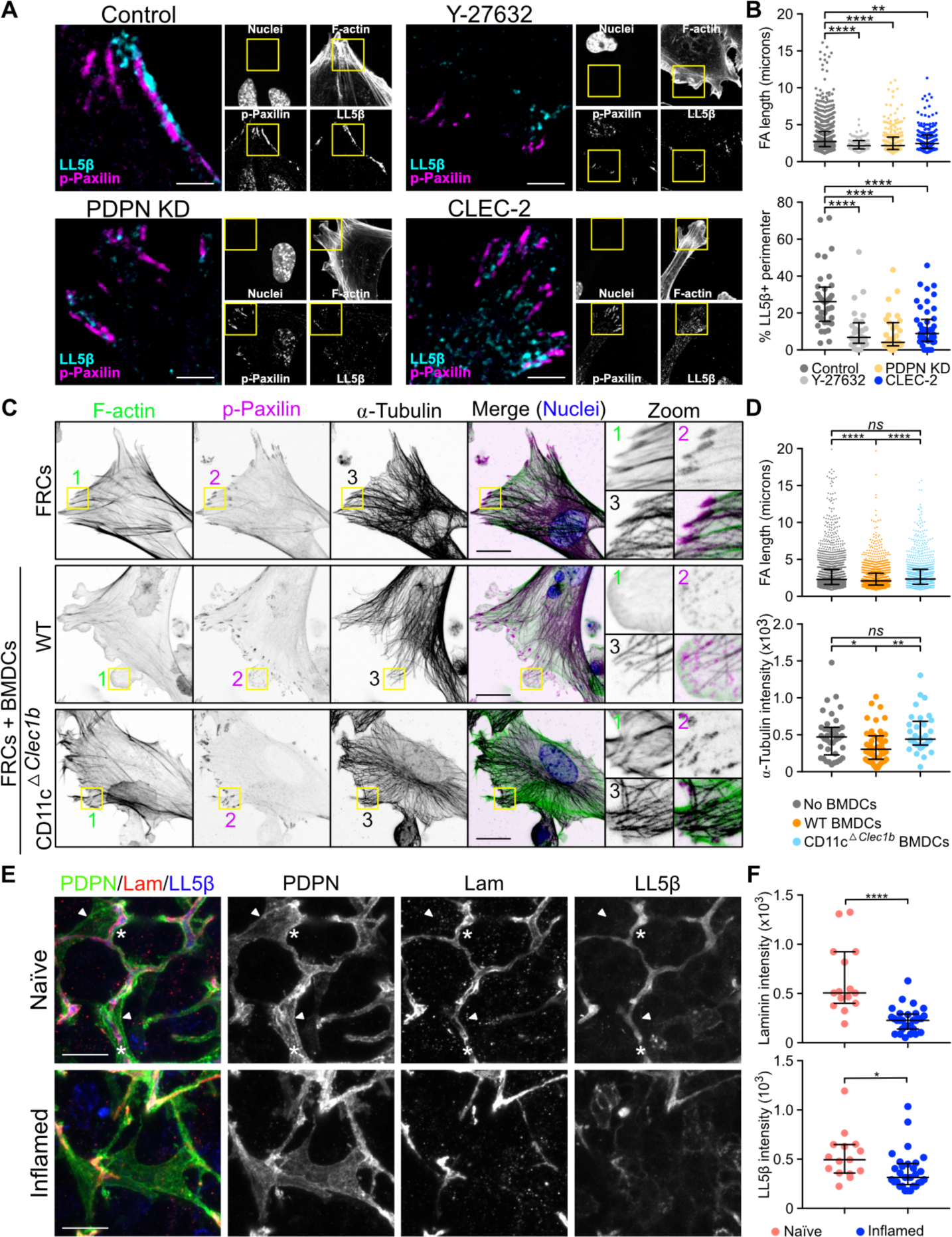
Focal adhesion, microtubule organization and contractility in FRCs. A-B) Immunofluorescence of FCR cell lines untreated, and control FRCs treated with Y-27632 ROCK inhibitor. A) Maximum z stack projections of representative images are shown. The scale bars represent 5 microns. B) Quantification of FA length from p-Paxilin staining and LL5*β* coverage as a percentage of total perimeter. Dots represent FAs (top graph) or single cells (bottom graph) (n=2). **P<0.005, ****P<0.00005, one-way ANOVA, Tukey’s multiple comparisons test. C) Immunofluorescence of three-dimensional cell culture of FRC plus wt and CD11cΔCLEC-2 BMDCs. Maximum z stack projections are shown. The scale bars rep-resent 20 microns. D) Quantification of FA length and *α*-tubulin intensity in the cortical area (10 microns deep) in FRCs. Dots represent FAs (top graph) or single cells (bottom graph) (n=3). *P<0.05, **P<0.005, ****P<0.00005, one-way ANOVA, Tukey’s multiple comparisons test. NS, not significant. E) Immunofluorescence of 20 microns thick cryosections of naïve and inflamed LNs from mice immunized with IFA/OVA. Maximum z stack projections are shown. The scale bars represent 20 microns. Asterisks and arrow heads indicate conduit-associated and conduit-independent surface of FRCs respectively. F) Quantification of the indicated conduit components within the PDPN network. Each dot represents the median grey intensity of a different region of interest (n=5). Error bars represent mean and SD. *P<0.05, ****P<0.00005, unpaired t test.

We next asked whether LL5*β* directs microtubule-mediated deposition of matrix components in the FRC network in vivo. High-resolution imaging of LN tissue revealed that in naïve LN, the entire FRC network expressed high levels of LL5*β*, and its localisation was always polarised towards the ensheathed conduit (Figure 5E). In contrast, in inflamed LNs, we observed many regions of the FRC network which lacked polarised localisation of LL5*β*, interestingly coinciding with loss of laminin in the same region (Figure 5E). This is a direct translation of the in vitro studies which predicted loss of LL5*β* when matrix adhesion is lost (Figure 4 and 5). These data require us to consider FRCs as polarised cells, exhibiting apical and basolateral polarity similarly to epithelial sheets (70) but enwrapping the conduit similarly to Schwann cells enwrapping nerve fibres (75). The inner surface of the FRC adheres to the conduit and recruits LL5*β* for ECM secretion, while the outer surface of the FRC excludes ECM, allowing optimal interaction with lymphocytes and antigen-presenting cells. Polarised and localised exocytosis in FRCs, directed by LL5*β*, can for the first time mechanistically explain how ECM components are exclusively found within the conduit and not elsewhere in the LN parenchyma.

### Changes in conduit flow and capture of antigen during LN expansion

We next asked how remodelling the ECM would affect conduit function. We compared the flow of 10, 70 and 500 kDa fluorescently-labelled dextrans through the LN sinuses and conduit network of naïve and acutely inflamed LNs. 10 kDa and 70 kDa dextrans rapidly percolated throughout LN parenchyma of naïve LNs and remained contained within the conduit, while the 500 kDa dextran remained within the capsule and was excluded from all but the most proximal branches of the conduit (Figure S4A). Similar results were found in inflamed LNs, meaning that the filtering function and size-exclusion of the conduit network is unaffected by LN expansion (Figure S4A). These results also indicated that ECM loss within parts of the network did not impede the overall flow of small soluble molecules through LNs.

However, looking closer at the pattern of the 10 kDa dextran-injected mice we noticed the presence of numerous gaps or interruptions in dextran flow throughout inflamed LNs (Figure 6A), which translated to a large decrease in percentage of dextran-positive area within the FRC network (Figure 6A, right panel). Interestingly, we found that dextran flow in inflamed LNs perfectly correlated with FRCs which had maintained polarised LL5*β* colocalised with basement membrane (laminin) (Figure 6B), indicating that conduit integrity determines conduit flow. Furthermore, despite the substantial reduction in conduit structures during acute expansion, FRCs maintain the overall global conduit functionality, reinforcing the robustness of the FRC network (76).

**Fig. 6.**
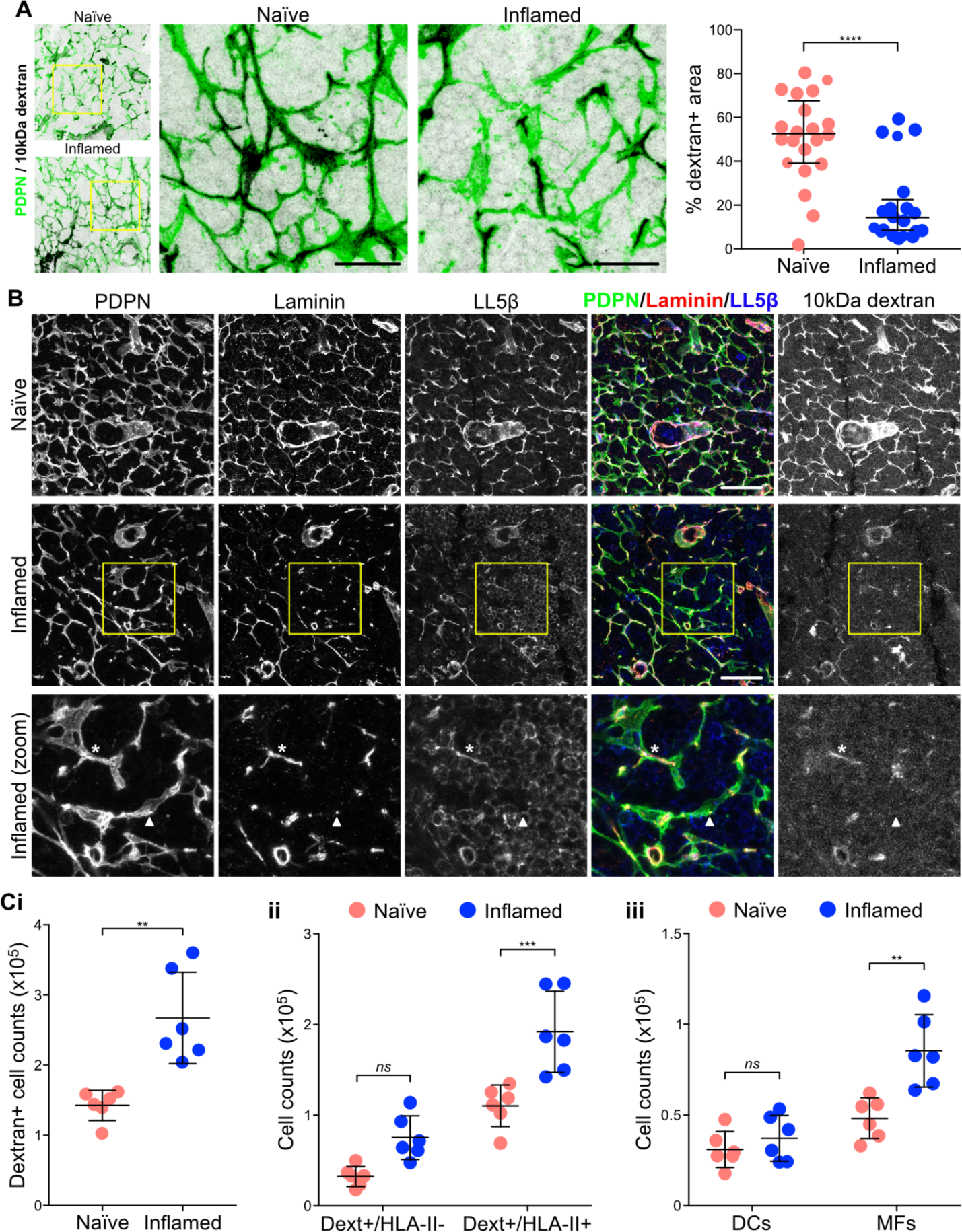
Conduit flow and antigen uptake in inflamed LNs. Mice were immunized by subcutaneous injection of IFA/OVA on the right flank. 5 days later, fluorescently labelled 10 kDa dextran was injected on both flanks. A) and B) Immunofluorescence of 20 microns thick cryosections of naïve and inflamed draining LNs 30 minutes post dextran injection. Maximum z stack projections are shown. The scale bars represent 20 microns (A), 40 microns (B) and 20 microns (B zoom). The asterisk indicates a portion of the FRC network with all conduit components plus dextrans flow. The arrow head indicates a portion of the FRC network where the conduit is not present and dextrans not flowing. Graph in A shows percentage of dextran-positive area within the PDPN-positive network. Each dot represents a different region of interest (n=6). Error bars represent mean and SD. ****P<0.00005, unpaired t test. Ci-iii) Flow cytometry analysis of draining LNs 90 minutes after dextran injection. The number of dextran-positive cells per LN within different cell subsets is shown. Each dot represents an individual (n=6). **P<0.005, ***P<0.0005, unpaired t test.

Antigen flow through the conduit secures under-control antigen sensing by LN resident cells (14–17, 77). We therefore questioned whether the interruptions in conduit flow ob-served in inflamed LNs would reflect leakiness of the system and whether this might affect antigen uptake by LN cells. By flow cytometry we found a significant increase in the number cells that captured the soluble 10 kDa dextran in inflamed LNs (Figure 6Ci and S5). This analysis was performed 90 minutes after dextran injection, meaning that antigen capture at the site of injection did not contribute to the effect. We observed a similar trend for the 70 kDa dextran, but no effect on the 500 kDa, confirming that size exclusion by the conduit is maintained during LN expansion (Figure S4B). It also indicates that the increase in antigen uptake is conduit-dependent, and not the result of more tracers being endocytosed from the subcapsular sinus. We found increased number of dextran+ cells within both HLAII- and HLA-II+ populations (Figure 6Cii), with HLA-II+ cells representing 80% of all dextran+ cells (Figure S5B). Previous studies have shown that CD11c+CD11b-/+ T-cell area DCs have privileged access to conduit content (15, 16), which could explain the similar numbers of dextran+ DCs we found in naïve and inflamed LNs (Figure 6Ciii). On the other hand, CD11b+CD11c- macrophages do not normally interact with the conduit (16), and nevertheless the number of dextran+ macrophages doubled in inflamed LNs (Figure 6Ciii). These changes in cell numbers resulted in underrepresentation of DCs within the dextran+HLA-II+ cell population (Figure S5B). Overall, these data further support that the conduit network becomes leaky during LN expansion, granting LN cells access to lymph-borne small molecules despite not being interacting with the conduit.

## Conclusions

Our results show for the first time that the integrity of the T-cell area conduit network is compromised during LN acute expansion and that ECM components are lost or effectively diluted during the early LN expansion. We observe discontinuous conduit flow within the FRC network, indicating areas of potential leakage. Increased number of dextran+/MHC-II+ cells in inflamed LNs support the notion of leaking conduits, increasing availability of soluble molecules during this phase of the immune response. We found that CLEC-2 binding regulates matrix remodelling by FRCs which occurs at both gene and protein levels and additionally downstream microtubule polarisation. CLEC-2 binding impairs focal adhesion stability and LL5*β* recruitment, temporarily inhibiting matrix deposition.

Previous works have described the complex architecture of the conduit network in steady state, and it is known that FRCs produce and organise ECM components (16–19). We show for the first time that FRCs exhibit apical/basolateral polarity and use polarised microtubule networks to direct ECM deposition unilaterally into conduit structures. Recruitment of LL5*β* facilitates docking and attachment of plus ends of microtubules to the cell membrane at sites of FRC-matrix adhesion, and enables matrix deposition. We previously proved that CLEC-2+ DCs inhibit PDPN-dependent contractility in FRCs (9), which also weakens FRC adhesion to the conduit. This in turn inhibits the recruitment of LL5*β* and FRC matrix secretion. LL5*β* expression is essential for FRCs to organise microtubules and form ECM matrices, and is exclusively localised basolaterally in association with laminin in vivo. Moreover, LL5*β* basolateral localisation is diminished during LN expansion.

Our results confirm that the FRC network remains connected in inflamed LNs (9, 10), even though the conduit flow is locally disrupted. Indicating that cell-cell connectivity is prioritised over and above maintaining ECM production. Our data suggest that to expand the LN rapidly, FRCs detach temporarily from the conduit, and halt matrix production, leading to loss of conduit integrity as pre-existing ECM fibres are stretched. However, the remaining intact sections of conduit are sufficient to channel the lymph throughout the LN parenchyma. Recent observations based on FRC ablation and a graph theory-based systems biology approach have demonstrated that FRCs establish “small-world” networks (76, 78). High local connectivity ensures the high topological robustness of the FRC network, which can tolerate loss of 50% of FRCs (76). We propose that a similar principle applies to conduit flow, in which interruption of the conduit network is efficiently overcome by sufficient alternative routes.

Conduit size exclusion depends on PLVAP expression by the lymphatic endothelial cells lining the sinus (79). Our data provides evidence that this barrier remains intact during early LN expansion. However, within the parenchyma, conduit leakiness enhances access of LN macrophages to small molecules. It is tempting to speculate that increased free diffusion of these draining molecules may influence the acute inflammatory response in LNs, providing a fast route for peripheral clues to prime LN cell populations.

In contexts outside of the LN, PDPN is often upregulated by fibroblasts in inflammatory settings (80–82). In these scenarios, the ECM deposition by these fibroblasts may also be regulated and modified by contact with CLEC-2+ myeloid cells or platelets leaking from inflamed vessels. It will be interesting to understand if the same CLEC-2-dependent transcriptional and protein expression regulation occurs in other PDPN+ fibroblasts. The CLEC-2/PDPN signalling axis is also a strong regulator of contractility in FRCs so this mechanism may also translate to how ECM is aligned and organised in other tissues.

## Experimental Proceedures

### Mice

Experiments were performed in accordance with national and institutional guidelines for animal care and approved by the Institutional Animal Ethics Committee Review Board, Cancer Research UK and the UK Home Office. Wild-type C57BL/6J mice were purchased from Charles River Laboratories. PDGFRaKI-H2BGFP mice (B6.129S4-Pdgfratm11(EGFP)Sor/J) were purchased from Jackson Laboratories. Generation of CD11cΔCLEC-2 was achieved as previously described (9) by crossing Clec1bfl/fl with CD11c-Cre mice (B6.Cg-Tg(Itgax-cre)1.1Reiz; gift from B. Reizis). Both males and females were used for in vivo and in vitro experiments and were aged 8–12 weeks. Cre-negative littermates were used as controls in all experiments.

### Tissue clearing and immunostaining of intact LNs

We used a modified version of the PACT (passive clarity technique) for whole LN staining based on previous publication (26). In brief, AntigenFix (DiaPath) fixed LNs were incubated overnight at 4°C in 40% acrylamide + 25 mg/ml 2,2-Azobis(2-methylpropionamidine) dihydrochloride (Sigma). Infused samples were degassed with nitrogen for 5 min and then incubated for 6 hr at 37°C. Samples were washed in PBS for 24 hr and incubated for 4 days with 8% SDS PBS solution at 37°C. After washing with PBS for 24 hr, LNs were incubated with 1:100 anti-collagen IV (table 1) in PBS 2% goat serum (Sigma-Aldrich) 0.1% triton X-100 (Sigma-Aldrich) 0.01% sodium azide (Webscientific) for 4 days at 37°C (rotation). Same concentration of the antibody was added after 1 and 2 days. Then wash in PBS at 37°C (rotation) for 24 hr and incubated with the secondary antibody in same conditions as the primary. Samples were transferred to a 2 g/ml Histodenz (Sigma-Aldricht), 0.01% sodium azide solution in PBS, incubated for 24 hr and imaged in this medium. Imaging was performed on a Leica TCS SP8 STED 3X using HC FLUOTAR L VISIR 25x water lenses.

### Electron microscopy of LN conduits

LNs were fixed overnight in 2%PFA/1.5% glutaraldehyde (both EM grade from TAAB) in 0.1M sodium Cacodylate at 4°C and embedded in 2.8% low melting point agarose dissolved in PBS. Slices of 100µm thickness were then cut in cold PBS using a vibrating microtome (VT1200S; Leica) and returned to fresh fix solution for a further 15 mins. Slices were then secondarily fixed for 1 h in 1% osmium tetraoxide/1.5% potassium ferricyanide at 4°C and then treated with 1% tannic acid in 0.1M sodium cacodylate for 45min at room temperature. Samples were then dehydrated in sequentially increasing concentration of ethanol solutions, and embedded in Epon resin. The 70nm ultrathin resin sections were cut with a Diatome 45° diamond knife using an ultramicrotome (UC7; Leica). Sections were collected on 1 ×2mm formvar-coated slot grids and stained with Reynolds lead citrate. All samples were imaged using a transmission electron microscope (Tecnai T12; FEI) equipped with a charge-coupled device camera (SIS Morada; Olympus).

### Immunisations

Mice were immunized via subcutaneous injection in the right flank of 100 µl of an emulsion of OVA in CFA or IFA (100 µg OVA per mouse) (Hooke Laboratories). After 5 days, mice were culled and inguinal LNs from both flanks (naïve and inflamed) were extracted for paired histological studies or flow cytometry analysis.

### Immunostaining of tissue sections

LN samples were fixed in AntigenFix (DiaPath) overnight, washed and incubated in PBS 30% sucrose (w/v) (Sigma-Aldrich) overnight at 4°C. Samples were embedded in Tissue-Tek^®^ O.C.T. Compound (Thomas Scientific) and frozen using 2-Methylbutane (GPR RECTAPUR), cooled with liquid nitrogen. 20 µm sections were cut using a Leica CM1850 cryostat. For immunostaining, tissue sections were blocked for 2 hr at room temperature in 10% goat normal Serum (Sigma-Aldrich), 0.3% Triton X-100 (Sigma-Aldrich) in PBS. Primary antibodies (table 1) were incubated overnight at 4°C in 10% goat normal Serum (Sigma-Aldrich), 0.01% Triton X-100 (Sigma-Aldrich) in PBS. 3 washing steps were used to remove unbound antibody, before incubation with the secondary antibody plus Hoechst (Fisher Scientific) for 2 hr at room temperature. Samples were washed and mounted in mowiol. Samples were imaged on a Leica TCS SP8 STED 3X using HC PL APO CS2 /1.4 63x oil lenses.

### Image analysis

Conduit components: Podoplanin staining was used to define the FRC network in LN frozen tissue sections. Podoplanin signal was filtered by Gaussian Blur (sigma=2) to remove background and thresholded identically in all samples. We next created a selection that was used in the corresponding channels in order to obtain the median intensity of the conduit components. We performed this process in a number of regions of interest within the T-cell area for each LN, always minimising the presence of vasculature. Semiautomated quantification of focal adhesion length: Signal for phospho-Paxilin staining was thresholded equally in all samples after removal of background noise by Gaussian Blur (sigma=2). Focal adhesions were segmented using the analyse particle tool in Fiji and fit ellipse. Major axis of the ellipse was used as an estimation of focal adhesion length.

### Cell culture

Cell lines: In vitro experiments were performed using inmortalized WT (control) and PDPN knockdown mouse FRC cell lines previously described (9). For CLEC-2-Fc expression by FRCs, Clec1b cDNA was cloned into pFUSE-rIgG-Fc2 plasmids (Invivogen) and transfected into WT FRCs using lipofectamine 2000 (Thermo Fisher Scientific). Transfected cells were selected by prolonged culture with zeocin 100 µg/ml (Invivogen) and secretion of CLEC-2-Fc was confirmed by western blotting for cell-derived supernatants (data not shown). FRC cell lines were cultured in DMEM plus glutamax (Life Technologies, Invitrogen) supplemented with 10% FBS, Penicillin-Streptomycin (100 U/mL) and 1% Insulin-Transferrin-Selenium (Life Technologies, Invitrogen) at 37°C in 10% CO2. Cells were passaged when they reach 80-90% confluence, by incubating in Cell Dissociation Buffer (Thermo Fisher Scientific) for 10 minutes at 37°C, plus a gentle treatment of 1 min with Trypsin 0.25% (Thermo Fisher Scientific). When indicated, FRCs were treated with 50 µg/ml CLEC-2-Fc or 10 µM ROCK inhibitor Y-27632 dihydrochloride (tocris) for the last 2 hr of culture. Primary cultures: Bone marrow cells were obtained from tibias and femurs from CD11cΔCLEC-2 mice and Cre-negative control littermates. Whole bone marrow was cultured in non-treated 10 cm petri dishes in RPMI media supplemented with 10% FBS and Penicillin-Streptomycin (100 U/mL) plus 20 ng/ml of recombinant murine GM-CSF (Pe-protech) at 4×106 cells / 13 ml of medium. After 3 days, cultures were supplemented with 4 ml of fresh media plus 37.2 ng/ml GM-CSF. After 6 days in culture, BMDCs were stimulated with 10 ng/ml Lipopolysaccharides from Escherichia coli 0111:B4 (Sigma-Aldrich) for 24 hr before harvesting.

### RNAseq analysis

FRCs were cultured for 24h, adding 50 µg/ml of CLEC-2-Fc from then beginning or 6 hours before collecting cells. Cells were left untreated as a control. RNA extractions were performed using the RNAeasy kit (Qiagen) following the manufacturer’s instructions, including a DNA digestion step to avoid genome contamination in further analysis. For transcriptome sequencing and analysis, RNA preparations from FRCs were sequenced to a depth of 9 million to 22 million reads by Queen Mary University (QMUL) Genome Centre. The raw read quality control was performed by the QMUL Genome Centre using Basesapce Illumina software. Paired end FASTQ files were then aligned to mus musculus GRCm38 reference assembly using STAR aligner software (83). Transcripts were assembled and relative transcript abundance were calculated using Salmon software (84). Using R (v3.4.4) and the Bioconductor tximport package (85), TPM (Transcripts per million) values were generated and annotated with ENSEMBL gene IDs. Bulk TPM data were categorised by the fold change (>2 fold) between control, 6 hr and 24 hr conditions using an in-house developed R script. Gene Ontology analysis were performed using the PANTHER software (28, 29) and PCA plots were generated using the ggplot package in R.

### FRC-derived matrices

FRC-derived matrices were generated in vitro according to published methods (86). In brief, gelatin-coated wells were used to culture FRC cell lines at 5×103 cells/cm2 in culture media supplemented with 50 µg/ml L(+)-Ascorbic acid sodium salt (Sigma-Aldrich) for 5 days, unless otherwise stated. Supplemented media was replenished at day 1 and 3. For proteomic analysis, cells in their matrix were collected in PBS, centrifuged and resuspended in 4M Urea. For microscopy analysis, cells were lysed incubating for 15 min at 37°C in PBS 1% Triton X-100 20 mM ammonium hydroxide. Matrices were blocked with 2% bovine serum albumin (w/v) (Sigma-Aldrich) and stained with the indicated antibodies (table 1). Samples were imaged with a Leica TCS SP5 Confocal Microscope using 63X oil HCX PL APO lenses.

### Proteomics of FRC-derived matrices

Quantitative proteomic analysis of the FRC-derived matrices was performed by sequential window acquisition of all theoretical spectra mass spectrometry (SWATH MS). For construction of the spectral library, FRCs and derived matrices were washed in PBS, centrifuged and enriched for extracellular matrix as previously described in Krasny et al. (2018). Enriched matrices were digested using gel-assisted protocol Shevchenko:2006co and desalted prior analysis by liquid chromatography-tandem mass spectrometry (LC-MS/MS) on Agilent 1260 HPLC coupled to TripleTOF 5600+ (SCIEX) mass spectrometer in data-dependent acquisition mode. For LC-MS/MS, peptides were spiked with iRT peptides (Biognosys AG), loaded on a 75 µm x 15 cm long analytical column packed with Reprosil Pur C18-AQ 3 µm resin (Dr. Maisch) end eluted using a linear gradient of 2-40% of Buffer B (98% ACN, 0.1% FA) in 90 min at flow rate of 250nl/min. Acquired datasets were searched by ProteinPilot 5.0.1 software (Sciex) against a Swissprot mouse database and spectral library was generated in Spectronaut 11 (Biognosys AG) from the results and combined with previously published library (Krasny et al. 2018). For quantitative analysis, FRCs and derived matrices were lysed on ice in 8M Urea, 100mM ammonium bicarbonate buffer and digested using gel-assisted protocol. Desalted peptides were spiked with iRT peptides and analysed on the same LC-MS/MS instrument using identical LC conditions. MS/MS data were acquired in 60 SWATH windows with fixed size of 13 Da. SWATH spectra were analysed in Spectronaut 11 with FDR restricted to 1%. Further statistical processing of median normalized data was performed in Perseus (1.5.6, ref Tyanova et al. Nat. Methods, 2016)

### Isobaric Tandem Mass Tag (TMT) Phosphoproteomics

Isobaric Tandem Mass Tag (TMT) Phosphoproteomics were performed as described in Tape et al., Cell, 2016 (PMID: 27087446). Following treatment, FRCs were lysed in 6 M urea, 10 mM NaPPi, 20 mM HEPES, pH 8.0, sonicated, centrifuged to clear cell debris, and protein concentration was determined by BCA (Pierce 23225). 200 µg of each condition were individually digested by FASP (87), amine-TMT-10-plex-labeled (Pierce 90111) on membrane (iFASP) (88), eluted, pooled, lyophilized, and subjected to automated phosphopeptide enrichment (APE) (89). Phosphopeptides were desalted using OLIGO R3 resin (Life Technologies 1-1339-03) and lyophilized prior to liquid chromatography-tandem mass spectrometry (LC-MS/MS) analysis. Samples were run on a Q-Exactive Plus mass spectrometer (Thermo Scientific) coupled to a Dionex Ultimate 3000 RSLC nano system (Thermo Scientific). Reversed-phase chromatographic separation was performed on a C18 PepMap 300 Å trap cartridge (0.3 mm i.d. x 5 mm, 5 µm bead size; loaded in a bi-directional manner), a 75 µm i.d. × 50 cm column (5 µm bead size) using a 120 minute linear gradient of 0-50% solvent B (MeCN 100% + 0.1% formic acid (FA)) against solvent A (H2O 100% + 0.1% FA) with a flow rate of 300 nL/min. The mass spectrometer was operated in the data-dependent mode to automatically switch between Orbitrap MS and MS/MS acquisition. Survey full scan MS spectra (from m/z 400-2000) were acquired in the Orbitrap with a resolution of 70,000 at m/z 400 and FT target value of 1×106 ions. The 20 most abundant ions were selected for fragmentation using higher-energy collisional dissociation (HCD) and dynamically excluded for 30 seconds. Fragmented ions were scanned in the Orbitrap at a resolution of 35,000 (TMT) at m/z 400. The isolation window was reduced to 1.2 m/z and a MS/MS fixed first mass of 120 m/z was used to aid TMT detection. For accurate mass measurement, the lock mass option was enabled using the polydimethylcyclosiloxane ion (m/z 445.120025) as an internal calibrant. For peptide identification, raw data files produced in Xcalibur 2.1 (Thermo Scientific) were processed in Proteome Discoverer 1.4 (Thermo Scientific) and searched against SwissProt mouse (2011-03 release, 15,082,690 entries) database using Mascot (v2.2). Searches were performed with a precursor mass tolerance set to 10 ppm, fragment mass tolerance set to 0.05 Da and a maximum number of missed cleavages set to 2. Peptides were further filtered using a mascot significance threshold <0.05, a peptide ion Score >20 and a FDR <0.01 (evaluated by Percolator (90). Phospho-site localization probabilities were calculated with phosphoRS 3.1 (>75%, maximum 4-PTM/peptide) (91). Phosphopeptides from Proteome Discoverer 1.4 were normalized against total protein levels (from in-gel digest experiments), and protein-level phospho-site locations (phosphoRS 3.1 score >75%, maximum 4-PTM/peptide) were manually annotated using PhosphoSitePlus. Phosphoproteomic volcano plots display mean Proteome Discoverer 1.4 quantification fold-difference values across all replicates (log2) against two-tailed t-test P values (calculated from arrays of raw MS/MS TMT intensity counts). Volcano plots were assembled in GraphPad Prism 6 (non-linear Gaussian regression, least squares fit). For principle component analysis (PCA), Proteome Discoverer 1.4 quantification ratio values were converted to log2, imported into R (version 3.0.1), computed using the function ‘princomp(X)’ and plotted in GraphPad Prism. Empirical parent kinases were manually identified by referenced Uniprot annotation and putative parent kinases were manually assigned using ScanSite (92) 3 (‘High-Stringency’ setting, top 0.2% of all sites, lowest score). Phospho-sites that did not meet these conditions were not annotated.

### GFP secretion assay

FRC cells lines were transfected with 500 ng of lumGFP plasmid (93) using Attractene Transfection Reagent (Qiagen) for 8 hr. Culture media was replenished with fresh media. After 15 hr, supernatants were collected and cells lysed in PBS 0.5% triton X-100. Supernatants were centrifuged in order to remove cell debris. GFP levels in cell lysates and supernatants were measured by a solid-phase sandwich ELISA (94). Briefly, polystyrene 96-well plates were coated overnight with 200 µl/well of PBS plus sheep anti-GFP 1:50,000 for 1 hr at room temperature. The antibody solution was removed and the plates were then incubated for 1 hr at room temperature to block nonspecific binding by using 300 µl/well of TEB (1% Triton X-100, 0.2% gelatin, 1 mM EDTA in PBS). The TEB was removed, each well was filled with 200 µl of samples or standard curve in TBE, and the plates were incubated while shaking for 1 hr. After extensive washing, plates were incubated with 200 µl TBE plus Rabbit anti-GFP 1:20,000 for 1 hr with shaking. Next, plates were washed and incubated with 200 µl/well of Goat anti-Rabbit HRP 1:3,000 in TBE for 1 hr plus shaking. Plates were washed three times in TBE and 3 times in PBS and using a standard o-phenylenediamine assay. Percentage of secreted GFP was calculated with respect to total GFP produced (supernatant plus lysates).

### Western Blotting

Equal number of cells were seeded and cultured for 24 hours. Cells were washed with cold PBS and lysed using Laemmli buffer (BioRad). All lysates were sonicated, heated for 10 min at 95°C and treated with 143 µM b-mercaptoethanol. Electrophoresis gels were loaded with the same quantity of lysates and run for 45 min at 130 V. Transfer to PDVF membranes were carried out at 65 V for 2 hr. Membranes were blocked for 2 hr at room temperature with 5 % skim milk powder (Sigma-Aldrich), 2 % BSA in PBS and stained with primary antibodies (table 1) overnight at 4°C in 1:5 diluted blocking buffer. The next day, membranes were thoroughly washed in PBS 0.05% Tween 20 and incubated with HRP-conjugated secondary antibodies 1:5000 in 1:5 diluted blocking buffer.

### Quantitative RT-PCR analysis for Phldb2 (LL5*β*) messenger RNA

cDNA was generated from RNA samples using the SuperScript™ IV Reverse Transcriptase kit (Thermo Fisher Scientific), following manufacturer’s instructions. Quantitative PCRs were run using the MESA Blue qPCR Mastermix (SYBR Assay). We used specific primers for detection of Phldb2 mRNA transcripts 1 and 2 (PrimerBank ID 23510303a1): Forward Primer AGC-CGCGTTTCTGAAAGCA (1653-1671); Reverse Primer CATCCGGGCGTCTTCCATT (1773-1755). Detection of GAPDH mRNA was used for normalization.

### Three-dimensional cell culture

FRCs were plated in 24well MatTek plates at 3.5×103 cell/cm2. Matured BMDCs were harvested and 150,000 cells were seeded per well in 150 µl collagen/matrigel matrix plus 20 ng/ml rmGM-CSF (95–97). FRC:BMDC ratio 1:43. Gels were set at 37°C for 30 min. After 24 hr, cells were fixed, permeabilized and stained for the stated cellular components (table 1).

### Immunostaining of cells in vitro

Cells were plated on 13 mm coverslips. Cells were fixed for 15 min in 3.6% formaldehyde and permeabilized with triton X-100 0.3% for 15 min, at room temperature. Cells were blocked with PBS 2% BSA for 1 hr at room temperature, followed by overnight incubation with corresponding primary antibodies (table 1) in PBS 1% BSA. After washing, cells were incubated with Alexa-fluor-conjugated secondary antibodies plus Hoechst and/or phalloidin to reveal DNA in cell nuclei and F-actin respectively, all in PBS 1% BSA for 2 hr at room temperature, washed and mounted on glass slides for imaging. Samples were imaged in Leica TCS SP5 and SP8 STED 3X Confocal Microscopes using 63X HCX PL APO lenses or HC PL APO CS2 /1.4 63x oil lenses.

### LL5*β* silencing by siRNA

WT FRC cell lines were transfected with four different siRNAs targeting LL5*β* expression (Dharmacon, GE Healthcare) using lipofectamine 2000 (Thermo Fisher Scientific). After 24 hr transfection, cells were washed and cultured in fresh media for an additional 12 hours before silencing efficiency was determined by qPCR. The following two siRNAs were selected for further assays: 1 GCAGAGUAUCAGCGGAACA and 2 GAACAAUGAAG-GACCGAGA. Scrambled RNA and non-template controls were used for comparison.

### Dextran uptake in vivo

Five days after immunization, mice were injected subcutaneously in both flanks with 20 µl of dextran solution (100 µg dextran per flank) conjugated to: Cascade Blue (10kDa dextran), Tetramethylrhodamine (70 kDa dextran) or Fluorescein (500 kDa dextran), all from Thermo Fisher Scientific. Mice were culled and paired inguinal LNs (inflamed vs non-inflamed) collected after 30 or 90 minutes for histological and flow cytometry analysis respectively.

### Flow cytometry of LNs

Inguinal LNs were carefully dissected and digested using collagenase P at 200 µg/ml (Sigma-Aldrich), dispase II 800 µg/ml (thermos fisher scientific) and DNase I 100 µg/ml (Sigma-Aldrich) in RPMI at 37°C in a water bath. Every 10 min LNs were mixed by pipetting up and down and half of the digestion media replenished by fresh until all tissue was digested. Cell suspensions were centrifuged, resuspended in FACS buffer (PBS 2 % FBS 10mM EDTA) and filtered through a 70 µm cell strainer (Corning). Cells were counted and approximately 1×106 cells were used for immunofluorescence staining. In brief, cells were resuspended in 100 µl FACS buffer, treated with Fc blocking for 10 min on ice and incubated with the indicated antibodies (table 1) for 30 min on ice. Cells were washed extensively and resuspended in 500 µl FACS buffer. Precision Count Beads (BioLegend) were used for accurate cell count. Samples were run in a Fortessa X20 flow cytometer (BD Biosciences) at the UCL Cancer Institute and analysed using the FlowJo software (FlowJo, LLC). Live cells were gated by FSC/SSC parameters and doublets discriminated by comparing SSC-A versus SSC-H.

### Statistical analysis

Statistical analysis was performed using Prism 7 (GraphPad Software). For in vivo experiments, naïve versus inflamed LNs were compared by unpaired, parametric t test, assuming that both populations had the comparable standard deviation. For in vitro experiments and all other multiple comparisons, ordinary one-way ANOVA followed by Tukey’s multiple comparisons test was performed. Binomial test type for PANTHER Overrepresentation Test of cellular components (RNAseq) and biological process (Phos-phoproteomics) was used to analysed changes induced by CLEC-2 binding to FRCs.

## Supporting information

Supplementary Figures_Martinez et al.

Proteomics raw data

## ACKNOWLEDGEMENTS

The authors are grateful to J. Sanes for LL5 antibody, and to C. Reis e Sousa for CD11cCLEC-2 mice; A. Vaughan for assistance with microscopy, and C. Bennett for assistance with in vivo experiments. They are also grateful to C. Bennett, S. Makris, C.M. de Winde and E. Sahai for critical reading of the manuscript. This work is supported by a Cancer Research UK (CRUK-A19763) (to S.E.A) and Medical Research Council (MC_U12266B). C.T is supported by Cancer Research UK (C60693/A23783). T.S is supported by the Wellcome Trust (105604/Z/14/Z). S.D is supported by Swiss National Science Foundation (grants P2BSP3/158804 and P300PA_167657) and EHA Research Grant award granted by the European Hematology Association. L.K. and P.H. are supported by the Institute of Cancer Research and Breast Cancer Now (2014NovPR360).

## AUTHOR CONTRIBUTIONS

V.G.M. and S.E.A designed the study and wrote the manuscript. V.G.M A.B and V.P performed in vitro experiments. V.G.M and S.D performed in vivo experiments. L.K. and P.H. performed mass spectrometry of ECM. C.T designed and performed TMT-Proteomics experiments. I.W and J.B. performed E.M of conduit structures, T.S, H.H and J. KV performed bioinformatics and statistical analyses. All authors contributed to editing the manuscript.

## Bibliography

1. Matthew B Buechler and Shannon J Turley. A short field guide to fibroblast function in immunity. Seminars in immunology, 35:48–58, February 2018.

2. Mario Novkovic, Lucas Onder, Hung-Wei Cheng, Gennady Bocharov, and Burkhard Ludewig. Integrative Computational Modeling of the Lymph Node Stromal Cell Landscape. Frontiers in Immunology, 9:2428, 2018.

3. Lauren B Rodda, Erick Lu, Mariko L Bennett, Caroline L Sokol, Xiaoming Wang, Sanjiv A Luther, Ben A Barres, Andrew D Luster, Chun Jimmie Ye, and Jason G Cyster. Single-Cell RNA Sequencing of Lymph Node Stromal Cells Reveals Niche-Associated Heterogeneity. Immunity, 48(5):1014–1028.e6, May 2018.

4. Alexander Link, Tobias K Vogt, Stéphanie Favre, Mirjam R Britschgi, Hans Acha-Orbea, Boris Hinz, Jason G Cyster, and Sanjiv A Luther. Fibroblastic reticular cells in lymph nodes regulate the homeostasis of naive T cells. Nature Immunology, 8(11):1255–1265, November 2007.

5. Viviana Cremasco, Matthew C Woodruff, Lucas Onder, Jovana Cupovic, Janice M Nieves-Bonilla, Frank A Schildberg, Jonathan Chang, Floriana Cremasco, Christopher J Harvey, Kai Wucherpfennig, Burkhard Ludewig, Michael C Carroll, and Shannon J Turley. B cell homeostasis and follicle confines are governed by fibroblastic reticular cells. Nature Immunology, 15(10):973–981, October 2014.

6. Je-Wook Lee, Mathieu Epardaud, Jing Sun, Jessica E Becker, Alexander C Cheng, Airis Yonekura, Joan K Heath, and Shannon J Turley. Peripheral antigen display by lymph node stroma promotes T cell tolerance to intestinal self. Nature Immunology, 8(2):181–190, February 2007.

7. Anne L Fletcher, Veronika Lukacs-Kornek, Erika D Reynoso, Sophie E Pinner, Angelique Bellemare-Pelletier, Mark S Curry, Ai-Ris Collier, Richard L Boyd, and Shannon J Turley. Lymph node fibroblastic reticular cells directly present peripheral tissue antigen under steady-state and inflammatory conditions. The Journal of Experimental Medicine, 207(4): 689–697, April 2010.

8. Juan Dubrot, Fernanda V Duraes, Lambert Potin, Francesca Capotosti, Dale Brighouse, Tobias Suter, Salomé LeibundGut-Landmann, Natalio Garbi, Walter Reith, Melody A Swartz, and Stéphanie Hugues. Lymph node stromal cells acquire peptide-MHCII complexes from dendritic cells and induce antigen-specific CD4 T cell tolerance. The Journal of Experimental Medicine, 211(6):1153–1166, June 2014.

9. Sophie E Acton, Aaron J Farrugia, Jillian L Astarita, Diego Mourão-Sá, Robert P Jenkins, Emma Nye, Steven Hooper, Janneke van Blijswijk, Neil C Rogers, Kathryn J Snelgrove, Ian Rosewell, Luis F Moita, Gordon Stamp, Shannon J Turley, Erik Sahai, and Caetano Reis e Sousa. Dendritic cells control fibroblastic reticular network tension and lymph node expansion. Nature, 514(7523):498–502, October 2014.

10. Jillian L Astarita, Viviana Cremasco, Jianxin Fu, Max C Darnell, James R Peck, Janice M Nieves-Bonilla, Kai Song, Yuji Kondo, Matthew C Woodruff, Alvin Gogineni, Lucas Onder, Burkhard Ludewig, Robby M Weimer, Michael C Carroll, David J Mooney, Lijun Xia, and Shannon J Turley. The CLEC-2–podoplanin axis controls the contractility of fibroblastic reticular cells and lymph node microarchitecture. Nature Immunology, 16(1):75–84, October 2014.

11. Andrea J Radtke, Wolfgang Kastenmüller, Diego A Espinosa, Michael Y Gerner, Sze-Wah Tse, Photini Sinnis, Ronald N Germain, Fidel P Zavala, and Ian A Cockburn. Lymph-node resident CD8–+ dendritic cells capture antigens from migratory malaria sporozoites and induce CD8+ T cell responses. PLoS pathogens, 11(2):e1004637, February 2015.

12. Gwendalyn J Randolph, Stoyan Ivanov, Bernd H Zinselmeyer, and Joshua P Scallan. The Lymphatic System: Integral Roles in Immunity. Annual Review of Immunology, 35(1):31–52, April 2017.

13. Mirela Kuka and Matteo Iannacone. The role of lymph node sinus macrophages in host defense. Annals of the New York Academy of Sciences, 1319(1):38–46, June 2014.

14. J E Gretz, C C Norbury, A O Anderson, A E Proudfoot, and S Shaw. Lymph-borne chemokines and other low molecular weight molecules reach high endothelial venules via specialized conduits while a functional barrier limits access to the lymphocyte microenvironments in lymph node cortex. The Journal of Experimental Medicine, 192(10):1425–1440, November 2000.

15. Martijn A Nolte, Jeroen A M Beliën, Inge Schadee-Eestermans, Wendy Jansen, Wendy W J Unger, Nico Van Rooijen, Georg Kraal, and Reina E Mebius. A conduit system distributes chemokines and small blood-borne molecules through the splenic white pulp. The Journal of Experimental Medicine, 198(3):505–512, August 2003.

16. Michael Sixt, Nobuo Kanazawa, Manuel Selg, Thomas Samson, Gunnel Roos, Dieter P Reinhardt, Reinhard Pabst, Manfred B Lutz, and Lydia Sorokin. The conduit system transports soluble antigens from the afferent lymph to resident dendritic cells in the T cell area of the lymph node. Immunity, 22(1):19–29, January 2005.

17. Ramon Roozendaal, Thorsten R Mempel, Lisa A Pitcher, Santiago F Gonzalez, Admar Verschoor, Reina E Mebius, Ulrich H von Andrian, and Michael C Carroll. Conduits mediate transport of low-molecular-weight antigen to lymph node follicles. Immunity, 30(2):264–276, February 2009.

18. Gregg P Sobocinski, Katherine Toy, Walter F Bobrowski, Stephen Shaw, Arthur O Anderson, and Eric P Kaldjian. Ultrastructural localization of extracellular matrix proteins of the lymph node cortex: evidence supporting the reticular network as a pathway for lymphocyte migration. BMC immunology, 11(1):42, 2010.

19. Deepali Malhotra, Anne L Fletcher, Jillian Astarita, Veronika Lukacs-Kornek, Prakriti Tayalia, Santiago F Gonzalez, Kutlu G Elpek, Sook Kyung Chang, Konstantin Knoblich, Martin E Hemler, Michael B Brenner, Michael C Carroll, David J Mooney, Shannon J Turley, and Immunological Genome Project Consortium. Transcriptional profiling of stroma from inflamed and resting lymph nodes defines immunological hallmarks. Nature Immunology, 13(5):499–510, April 2012.

20. Meilang Xue and Christopher J Jackson. Extracellular Matrix Reorganization During Wound Healing and Its Impact on Abnormal Scarring. Advances in wound care, 4(3):119–136, March 2015.

21. Thomas A Wynn and Thirumalai R Ramalingam. Mechanisms of fibrosis: therapeutic translation for fibrotic disease. Nature Medicine, 18(7):1028–1040, July 2012.

22. Timothy W Schacker, Jason M Brenchley, Gregory J Beilman, Cavan Reilly, Stefan E Pambuccian, Jodie Taylor, David Skarda, Matthew Larson, Daniel C Douek, and Ashley T Haase. Lymphatic tissue fibrosis is associated with reduced numbers of naive CD4+ T cells in human immunodeficiency virus type 1 infection. Clinical and vaccine immunology: CVI, 13 (5):556–560, May 2006.

23. Ming Zeng, Anthony J Smith, Stephen W Wietgrefe, Peter J Southern, Timothy W Schacker, Cavan S Reilly, Jacob D Estes, Gregory F Burton, Guido Silvestri, Jeffrey D Lifson, John V Carlis, and Ashley T Haase. Cumulative mechanisms of lymphoid tissue fibrosis and T cell depletion in HIV-1 and SIV infections. The Journal of clinical investigation, 121(3):998–1008, March 2011.

24. Angela Riedel, David Shorthouse, Lisa Haas, Benjamin A Hall, and Jacqueline Shields. Tumor-induced stromal reprogramming drives lymph node transformation. Nature Immunology, 17(9):1118–1127, September 2016.

25. N A Rohner, J McClain, S L Tuell, A Warner, B Smith, Y Yun, A Mohan, M Sushnitha, and S N Thomas. Lymph node biophysical remodeling is associated with melanoma lymphatic drainage. The FASEB Journal, 29(11):4512–4522, November 2015.

26. Bin Yang, Jennifer B Treweek, Rajan P Kulkarni, Benjamin E Deverman, Chun-Kan Chen, Eric Lubeck, Sheel Shah, Long Cai, and Viviana Gradinaru. Single-cell phenotyping within transparent intact tissue through whole-body clearing. Cell, 158(4):945–958, August 2014.

27. Marc Bajénoff and Ronald N Germain. B-cell follicle development remodels the conduit system and allows soluble antigen delivery to follicular dendritic cells. Blood, 114(24):4989–4997, December 2009.

28. Huaiyu Mi, Xiaosong Huang, Anushya Muruganujan, Haiming Tang, Caitlin Mills, Diane Kang, and Paul D Thomas. PANTHER version 11: expanded annotation data from Gene Ontology and Reactome pathways, and data analysis tool enhancements. Nucleic acids research, 45(D1):D183–D189, January 2017.

29. Huaiyu Mi, Anushya Muruganujan, John T Casagrande, and Paul D Thomas. Large-scale gene function analysis with the PANTHER classification system. Nature protocols, 8(8): 1551–1566, August 2013.

30. Alexandra Naba, Karl R Clauser, Sebastian Hoersch, Hui Liu, Steven A Carr, and Richard O Hynes. The matrisome: in silico definition and in vivo characterization by proteomics of normal and tumor extracellular matrices. Molecular & cellular proteomics: MCP, 11(4): M111.014647, April 2012.

31. Alexandra Naba, Karl R Clauser, Huiming Ding, Charles A Whittaker, Steven A Carr, and Richard O Hynes. The extracellular matrix: Tools and insights for the “omics” era. Matrix Biology, 49:10–24, January 2016.

32. Alexandra Naba, Oliver M T Pearce, Amanda Del Rosario, Duanduan Ma, Huiming Ding, Vinothini Rajeeve, Pedro R Cutillas, Frances R Balkwill, and Richard O Hynes. Characterization of the Extracellular Matrix of Normal and Diseased Tissues Using Proteomics. Journal of proteome research, 16(8):3083–3091, August 2017.

33. Peter D Ellis, James C Metcalfe, Marko Hyvönen, and Paul R Kemp. Adhesion of endothelial cells to NOV is mediated by the integrins alphavbeta3 and alpha5beta1. Journal of vascular research, 40(3):234–243, May 2003.

34. Filippo Gagliardi, Ashwin Narayanan, and Pietro Mortini. SPARCL1 a novel player in cancer biology. Critical reviews in oncology/hematology, 109:63–68, January 2017.

35. Kai Yin, Mengyuan Shang, Shengchun Dang, Linjun Wang, Yiwen Xia, Lei Cui, Xin Fan, Jianguo Qu, Jixiang Chen, and Zekuan Xu. Netrin-1 induces the proliferation of gastric cancer cells via the ERK/MAPK signaling pathway and FAK activation. Oncology Reports, 40(4):2325–2333, August 2018.

36. Angara Sureshbabu, Hiroshi Okajima, Daisuke Yamanaka, Elizabeth Tonner, Surya Shastri, Joanna Maycock, Malgorzata Szymanowska, John Shand, Shin-Ichiro Takahashi, James Beattie, Gordon Allan, and David Flint. IGFBP5 induces cell adhesion, increases cell survival and inhibits cell migration in MCF-7 human breast cancer cells. Journal of Cell Science, 125(Pt 7):1693–1705, April 2012.

37. Koji Ohashi, Takashi Enomoto, Yusuke Joki, Rei Shibata, Yasuhiro Ogura, Yoshiyuki Kataoka, Yuuki Shimizu, Takahiro Kambara, Yusuke Uemura, Daisuke Yuasa, Kazuhiro Matsuo, Satoko Hayakawa, Mizuho Hiramatsu-Ito, Toyoaki Murohara, and Noriyuki Ouchi. Neuron-derived neurotrophic factor functions as a novel modulator that enhances endothelial cell function and revascularization processes. The Journal of biological chemistry, 289 (20):14132–14144, May 2014.

38. Wei Jia, Hong Li, and You-Wen He. The extracellular matrix protein mindin serves as an integrin ligand and is critical for inflammatory cell recruitment. Blood, 106(12):3854–3859, December 2005.

39. En-Lin Song, Ya-Ping Hou, Shen-Ping Yu, Sheng-Guo Chen, Jun-Ting Huang, Tao Luo, Ling-Ping Kong, Jie Xu, and Hua-Qiao Wang. EFEMP1 expression promotes angiogenesis and accelerates the growth of cervical cancer in vivo. Gynecologic oncology, 121(1):174–180, April 2011.

40. Susana de Vega, Kentaro Hozumi, Nobuharu Suzuki, Risa Nonaka, Eimi Seo, Anna Takeda, Tomoko Ikeuchi, Motoyoshi Nomizu, Yoshihiko Yamada, and Eri Arikawa-Hirasawa. Identification of peptides derived from the C-terminal domain of fibulin-7 active for endothelial cell adhesion and tube formation disruption. Biopolymers, 106(2):184–195, March 2016.

41. Chang H Lee, Bhranti Shah, Eduardo K Moioli, and Jeremy J Mao. CTGF directs fibroblast differentiation from human mesenchymal stem/stromal cells and defines connective tissue healing in a rodent injury model. The Journal of clinical investigation, 120(9):3340–3349, September 2010.

42. Daisuke Niimori, Rie Kawano, Kanako Niimori-Kita, Hironobu Ihn, and Kunimasa Ohta. Tsukushi is involved in the wound healing by regulating the expression of cytokines and growth factors. Journal of cell communication and signaling, 8(3):173–177, September 2014.

43. Mitsuaki Ono, Asuka Masaki, Azusa Maeda, Tina M Kilts, Emilio S Hara, Taishi Komori, Hai Pham, Takuo Kuboki, and Marian F Young. CCN4/WISP1 controls cutaneous wound healing by modulating proliferation, migration and ECM expression in dermal fibroblasts via –5—1 and TNF–. Matrix Biology, 68-69:533–546, August 2018.

44. Yasunori Enomoto, Sayomi Matsushima, Kiyoshi Shibata, Yoichiro Aoshima, Haruna Yagi, Shiori Meguro, Hideya Kawasaki, Isao Kosugi, Tomoyuki Fujisawa, Noriyuki Enomoto, Naoki Inui, Yutaro Nakamura, Takafumi Suda, and Toshihide Iwashita. LTBP2 is secreted from lung myofibroblasts and is a potential biomarker for idiopathic pulmonary fibrosis. Clinical science (London, England: 1979), 132(14):1565–1580, July 2018.

45. Luigi Mario Castello, Davide Raineri, Livia Salmi, Nausicaa Clemente, Rosanna Vaschetto, Marco Quaglia, Massimiliano Garzaro, Sergio Gentilli, Paolo Navalesi, Vincenzo Cantaluppi, Umberto Dianzani, Anna Aspesi, and Annalisa Chiocchetti. Osteopontin at the Crossroads of Inflammation and Tumor Progression. Mediators of Inflammation, 2017:4049098–22, 2017.

46. Devadarssen Murdamoothoo, Anja Schwenzer, Jessica Kant, Tristan Rupp, Anna Marzeda, Kim Midwood, and Gertraud Orend. Investigating cell-type specific functions of tenascin-C. Methods in cell biology, 143:401–428, 2018.

47. Zhi-Qin Wang, Wen-Ming Xing, Hua-Hua Fan, Ke-Sheng Wang, Hai-Kuo Zhang, Qin-Wan Wang, Jia Qi, Hong-Meng Yang, Jie Yang, Ya-Na Ren, Shu-Jian Cui, Xin Zhang, Feng Liu, Dao-Hong Lin, Wen-Hui Wang, Michael K Hoffmann, and Ze-Guang Han. The novel lipopolysaccharide-binding protein CRISPLD2 is a critical serum protein to regulate endotoxin function. Journal of immunology (Baltimore, Md.: 1950), 183(10):6646–6656, November 2009.

48. Axel Seher, Joachim Nickel, Thomas D Mueller, Susanne Kneitz, Susanne Gebhardt, Tobias Meyer ter Vehn, Guenther Schlunck, and Walter Sebald. Gene expression profiling of connective tissue growth factor (CTGF) stimulated primary human tenon fibroblasts reveals an inflammatory and wound healing response in vitro. Molecular vision, 17:53–62, January 2011.

49. Marwa M Qadri, Gregory D Jay, Rennolds S Ostrom, Ling X Zhang, and Khaled A Elsaid. cAMP attenuates TGF-—’s profibrotic responses in osteoarthritic synoviocytes: involvement of hyaluronan and PRG4. American journal of physiology. Cell physiology, 315(3):C432–C443, September 2018.

50. Nikolaos A Afratis, Dragana Nikitovic, Hinke A B Multhaupt, Achilleas D Theocharis, John R Couchman, and Nikos K Karamanos. Syndecans - key regulators of cell signaling and biological functions. The FEBS journal, 284(1):27–41, January 2017.

51. Mailin Julia Hamm, Bettina Carmen Kirchmaier, and Wiebke Herzog. Sema3d controls collective endothelial cell migration by distinct mechanisms via Nrp1 and PlxnD1. The Journal of cell biology, 215(3):415–430, November 2016.

52. Andrea Casazza, Pietro Fazzari, and Luca Tamagnone. Semaphorin signals in cell adhesion and cell migration: functional role and molecular mechanisms. Advances in experimental medicine and biology, 600(Chapter 8):90–108, 2007.

53. Stephane Esnault, Elizabeth E Torr, Ksenija Bernau, Mats W Johansson, Elizabeth A Kelly, Nathan Sandbo, and Nizar N Jarjour. Endogenous Semaphorin-7A Impedes Human Lung Fibroblast Differentiation. PloS one, 12(1):e0170207, 2017.

54. Panagiotis Flevaris and Douglas Vaughan. The Role of Plasminogen Activator Inhibitor Type-1 in Fibrosis. Seminars in thrombosis and hemostasis, 43(2):169–177, March 2017.

55. Huan Zhai, Xun Qi, Zixuan Li, Wei Zhang, Chenguang Li, Lu Ji, Ke Xu, and Hongshan Zhong. TIMP-3 suppresses the proliferation and migration of SMCs from the aortic neck of atherosclerotic AAA in rabbits, via decreased MMP-2 and MMP-9 activity, and reduced TNF-– expression. Molecular Medicine Reports, 18(2):2061–2067, August 2018.

56. Satoshi Nagamine, Michiko Tamba, Hisako Ishimine, Kota Araki, Kensuke Shiomi, Takuya Okada, Tatsuyuki Ohto, Satoshi Kunita, Satoru Takahashi, Ronnie G P Wismans, Toin H van Kuppevelt, Masayuki Masu, and Kazuko Keino-Masu. Organ-specific sulfation patterns of heparan sulfate generated by extracellular sulfatases Sulf1 and Sulf2 in mice. The Journal of biological chemistry, 287(12):9579–9590, March 2012.

57. E Kessler, K Takahara, L Biniaminov, M Brusel, and D S Greenspan. Bone morphogenetic protein-1: the type I procollagen C-proteinase. Science, 271(5247):360–362, January 1996.

58. Hosami Harada and Masaaki Takahashi. CD44-dependent intracellular and extracellular catabolism of hyaluronic acid by hyaluronidase-1 and-2. Journal of Biological Chemistry, 282(8):5597–5607, February 2007.

59. Sylvie Rodrigues-Ferreira, Mohamed Abdelkarim, Patricia Dillenburg-Pilla, Anny-Claude Luissint, Anne di Tommaso, Frédérique Deshayes, Carmen Lucia S Pontes, Angie Molina, Nicolas Cagnard, Franck Letourneur, Marina Morel, Rosana I Reis, Dulce E Casarini, Benoit Terris, Pierre-Olivier Couraud, Claudio M Costa-Neto, Mélanie Di Benedetto, and Clara Nahmias. Angiotensin II facilitates breast cancer cell migration and metastasis. PloS one, 7(4): e35667, 2012.

60. Jiraporn Tocharus, Akiho Tsuchiya, Miwa Kajikawa, Yoshifumi Ueta, Chio Oka, and Masashi Kawaichi. Developmentally regulated expression of mouse HtrA3 and its role as an inhibitor of TGF-beta signaling. Development, growth & differentiation, 46(3):257–274, June 2004.

61. Frederic Bost, Maryam Diarra-Mehrpour, and Jean-Pierre Martin. Inter-alpha-trypsin inhibitor proteoglycan family. A group of proteins binding and stabilizing the extracellular matrix. European Journal of Biochemistry, 252(3):339–346, March 1998.

62. Sarah Porter, Ian M Clark, Lara Kevorkian, and Dylan R Edwards. The ADAMTS metalloproteinases. The Biochemical journal, 386(Pt 1):15–27, February 2005.

63. Stephen P Evanko, Susan Potter-Perigo, Paul L Bollyky, Gerald T Nepom, and Thomas N Wight. Hyaluronan and versican in the control of human T-lymphocyte adhesion and migration. Matrix biology: journal of the International Society for Matrix Biology, 31(2):90–100, March 2012.

64. R Riessen, M Fenchel, H Chen, D I Axel, K R Karsch, and J Lawler. Cartilage oligomeric matrix protein (thrombospondin-5) is expressed by human vascular smooth muscle cells. Arteriosclerosis, thrombosis, and vascular biology, 21(1):47–54, January 2001.

65. Sawako Yoshina, Kenjiro Sakaki, Aki Yonezumi-Hayashi, Keiko Gengyo-Ando, Hideshi Inoue, Yuichi Iino, and Shohei Mitani. Identification of a novel ADAMTS9/GON-1 function for protein transport from the ER to the Golgi. Molecular Biology of the Cell, 23(9):1728–1741, May 2012.

66. Carolyn M Dancevic, Fiona W Fraser, Adam D Smith, Nicole Stupka, Alister C Ward, and Daniel R McCulloch. Biosynthesis and expression of a disintegrin-like and metalloproteinase domain with thrombospondin-1 repeats-15: a novel versican-cleaving proteoglycanase. The Journal of biological chemistry, 288(52):37267–37276, December 2013.

67. Robin Roychaudhuri, Anja H Hergrueter, Francesca Polverino, Maria E Laucho-Contreras, Kushagra Gupta, Niels Borregaard, and Caroline A Owen. ADAM9 is a novel product of polymorphonuclear neutrophils: regulation of expression and contributions to extracellular matrix protein degradation during acute lung injury. Journal of immunology (Baltimore, Md.: 1950), 193(5):2469–2482, September 2014.

68. Lee Roth, Rotem Kalev-Altman, Efrat Monsonego-Ornan, and Dalit Sela-Donenfeld. A new role of the membrane-type matrix metalloproteinase 16 (MMP16/MT3-MMP) in neural crest cell migration. The International journal of developmental biology, 61(3-4-5):245–256, 2017.

69. Christopher J Tape, Stephanie Ling, Maria Dimitriadi, Kelly M McMahon, Jonathan D Worboys, Hui Sun Leong, Ida C Norrie, Crispin J Miller, George Poulogiannis, Douglas A Lauffenburger, and Claus Jørgensen. Oncogenic KRAS Regulates Tumor Cell Signaling via Stromal Reciprocation. Cell, 165(7):1818, June 2016.

70. Ivar Noordstra and Anna Akhmanova. Linking cortical microtubule attachment and exocytosis. F1000Research, 6:469, 2017.

71. Gideon Lansbergen, Ilya Grigoriev, Yuko Mimori-Kiyosue, Toshihisa Ohtsuka, Susumu Higa, Isao Kitajima, Jeroen Demmers, Niels Galjart, Adriaan B Houtsmuller, Frank Grosveld, and Anna Akhmanova. CLASPs attach microtubule plus ends to the cell cortex through a complex with LL5beta. Developmental cell, 11(1):21–32, July 2006.

72. Azusa Hotta, Tomomi Kawakatsu, Tomoya Nakatani, Toshitaka Sato, Chiyuki Matsui, Taiko Sukezane, Tsuyoshi Akagi, Tomoko Hamaji, Ilya Grigoriev, Anna Akhmanova, Yoshimi Takai, and Yuko Mimori-Kiyosue. Laminin-based cell adhesion anchors microtubule plus ends to the epithelial cell basal cortex through LL5–/—. The Journal of cell biology, 189(5): 901–917, May 2010.

73. Samantha J Stehbens, Matthew Paszek, Hayley Pemble, Andreas Ettinger, Sarah Gierke, and Torsten Wittmann. CLASPs link focal-adhesion-associated microtubule capture to localized exocytosis and adhesion site turnover. Nature Cell Biology, 16(6):561–573, June 2014.

74. K Katoh, Y Kano, M Amano, K Kaibuchi, and K Fujiwara. Stress fiber organization regulated by MLCK and Rho-kinase in cultured human fibroblasts. American journal of physiology. Cell physiology, 280(6):C1669–79, June 2001.

75. Nicolas Tricaud. Myelinating Schwann Cell Polarity and Mechanically-Driven Myelin Sheath Elongation. Frontiers in cellular neuroscience, 11:414, 2017.

76. Mario Novkovic, Lucas Onder, Jovana Cupovic, Jun Abe, David Bomze, Viviana Cremasco, Elke Scandella, Jens V Stein, Gennady Bocharov, Shannon J Turley, and Burkhard Ludewig. Topological Small-World Organization of the Fibroblastic Reticular Cell Network Determines Lymph Node Functionality. PLoS biology, 14(7):e1002515, July 2016.

77. R T Palframan, S Jung, G Cheng, W Weninger, Y Luo, M Dorf, D R Littman, B J Rollins, H Zweerink, A Rot, and U H von Andrian. Inflammatory chemokine transport and presentation in HEV: a remote control mechanism for monocyte recruitment to lymph nodes in inflamed tissues. The Journal of Experimental Medicine, 194(9):1361–1373, November 2001.

78. Johannes Textor, Judith N Mandl, and Rob J de Boer. The Reticular Cell Network: A Robust Backbone for Immune Responses. PLoS biology, 14(10):e2000827, October 2016.

79. Pia Rantakari, Kaisa Auvinen, Norma Jäppinen, Maria Kapraali, Joona Valtonen, Marika Karikoski, Heidi Gerke, Imtiaz Iftakhar-E-Khuda, Johannes Keuschnigg, Eiji Umemoto, Kazuo Tohya, Masayuki Miyasaka, Kati Elima, Sirpa Jalkanen, and Marko Salmi. The endothelial protein PLVAP in lymphatics controls the entry of lymphocytes and antigens into lymph nodes. Nature Immunology, 16(4):386–396, April 2015.

80. Koji Shindo, Shinichi Aishima, Kenoki Ohuchida, Kenji Fujiwara, Minoru Fujino, Yusuke Mizuuchi, Masami Hattori, Kazuhiro Mizumoto, Masao Tanaka, and Yoshinao Oda. Podoplanin expression in cancer-associated fibroblasts enhances tumor progression of invasive ductal carcinoma of the pancreas. Molecular cancer, 12(1):168, December 2013.

81. Hyun-Yi Kim, Ki-Sang Rha, Geun Ae Shim, Ju-Hee Kim, Jin-Man Kim, Song Mei Huang, and Bon Seok Koo. Podoplanin is involved in the prognosis of head and neck squamous cell carcinoma through interaction with VEGF-C. Oncology Reports, 34(2):833–842, August 2015.

82. Kakeru Hisakane, Koichi Saruwatari, Satoshi Fujii, Keisuke Kirita, Shigeki Umemura, Shingo Matsumoto, Kiyotaka Yoh, Seiji Niho, Hironobu Ohmatsu, Takeshi Kuwata, Atsushi Ochiai, Akihiko Gemma, Masahiro Tsuboi, Koichi Goto, and Genichiro Ishii. Unique intravascular tumor microenvironment predicting recurrence of lung squamous cell carcinoma. Journal of cancer research and clinical oncology, 142(3):593–600, March 2016.

83. Alexander Dobin, Carrie A Davis, Felix Schlesinger, Jorg Drenkow, Chris Zaleski, Sonali Jha, Philippe Batut, Mark Chaisson, and Thomas R Gingeras. STAR: ultrafast universal RNA-seq aligner. Bioinformatics (Oxford, England), 29(1):15–21, January 2013.

84. Rob Patro, Geet Duggal, Michael I Love, Rafael A Irizarry, and Carl Kingsford. Salmon provides fast and bias-aware quantification of transcript expression. Nature methods, 14(4): 417–419, April 2017.

85. Charlotte Soneson, Michael I Love, and Mark D Robinson. Differential analyses for RNA-seq: transcript-level estimates improve gene-level inferences. F1000Research, 4:1521, 2015.

86. Janusz Franco-Barraza, Dorothy A Beacham, Michael D Amatangelo, and Edna Cukierman. Preparation of Extracellular Matrices Produced by Cultured and Primary Fibroblasts, volume 167 of Fibroblast Derived 3D Matrix. John Wiley & Sons, Inc., Hoboken, NJ, USA, May 2001.

87. Jacek R Wiśniewski, Alexandre Zougman, Nagarjuna Nagaraj, and Matthias Mann. Universal sample preparation method for proteome analysis. Nature methods, 6(5):359–362, May 2009.

88. Gary S McDowell, Aleksandr Gaun, and Hanno Steen. iFASP: combining isobaric mass tagging with filter-aided sample preparation. Journal of proteome research, 12(8):3809–3812, August 2013.

89. Christopher J Tape, Jonathan D Worboys, John Sinclair, Robert Gourlay, Janis Vogt, Kelly M McMahon, Matthias Trost, Douglas A Lauffenburger, Douglas J Lamont, and Claus Jørgensen. Reproducible automated phosphopeptide enrichment using magnetic TiO2 and Ti-IMAC. Analytical chemistry, 86(20):10296–10302, October 2014.

90. Lukas Käll, Jesse D Canterbury, Jason Weston, William Stafford Noble, and Michael J MacCoss. Semi-supervised learning for peptide identification from shotgun proteomics datasets. Nature methods, 4(11):923–925, November 2007.

91. Thomas Taus, Thomas Köcher, Peter Pichler, Carmen Paschke, Andreas Schmidt, Christoph Henrich, and Karl Mechtler. Universal and confident phosphorylation site localization using phosphoRS. Journal of proteome research, 10(12):5354–5362, December 2011.

92. John C Obenauer, Lewis C Cantley, and Michael B Yaffe. Scansite 2.0: Proteome-wide prediction of cell signaling interactions using short sequence motifs. Nucleic acids research, 31(13):3635–3641, July 2003.

93. R Blum, D J Stephens, and I Schulz. Lumenal targeted GFP, used as a marker of soluble cargo, visualises rapid ERGIC to Golgi traffic by a tubulo-vesicular network. Journal of Cell Science, 113 (Pt 18):3151–3159, September 2000.

94. Anastasia D Blagoveshchenskaya, Matthew J Hannah, Simon Allen, and Daniel F Cutler. Selective and signal-dependent recruitment of membrane proteins to secretory granules formed by heterologously expressed von Willebrand factor. Molecular Biology of the Cell, 13(5):1582–1593, May 2002.

95. Cedric Gaggioli, Steven Hooper, Cristina Hidalgo-Carcedo, Robert Grosse, John F Marshall, Kevin Harrington, and Erik Sahai. Fibroblast-led collective invasion of carcinoma cells with differing roles for RhoGTPases in leading and following cells. Nature Cell Biology, 9 (12):1392–1400, December 2007.

96. Fernando Calvo, Nil Ege, Araceli Grande-Garcia, Steven Hooper, Robert P Jenkins, Shahid I Chaudhry, Kevin Harrington, Peter Williamson, Emad Moeendarbary, Guillaume Charras, and Erik Sahai. Mechanotransduction and YAP-dependent matrix remodelling is required for the generation and maintenance of cancer-associated fibroblasts. Nature Cell Biology, 15(6):637–646, June 2013.

97. Sophie E Acton, Jillian L Astarita, Deepali Malhotra, Veronika Lukacs-Kornek, Bettina Franz, Paul R Hess, Zoltan Jakus, Michael Kuligowski, Anne L Fletcher, Kutlu G Elpek, Angelique Bellemare-Pelletier, Lindsay Sceats, Erika D Reynoso, Santiago F Gonzalez, Daniel B Graham, Jonathan Chang, Anneli Peters, Matthew Woodruff, Young-A Kim, Wojciech Swat, Takashi Morita, Vijay Kuchroo, Michael C Carroll, Mark L Kahn, Kai W Wucherpfennig, and Shannon J Turley. Podoplanin-Rich Stromal Networks Induce Dendritic Cell Motility via Activation of the C-type Lectin Receptor CLEC-2. Immunity, 37(2):276–289, August 2012.

