## Supplementary Figures_Martinez et al. for "Conduit integrity is compromised during acute lymph node expansion"

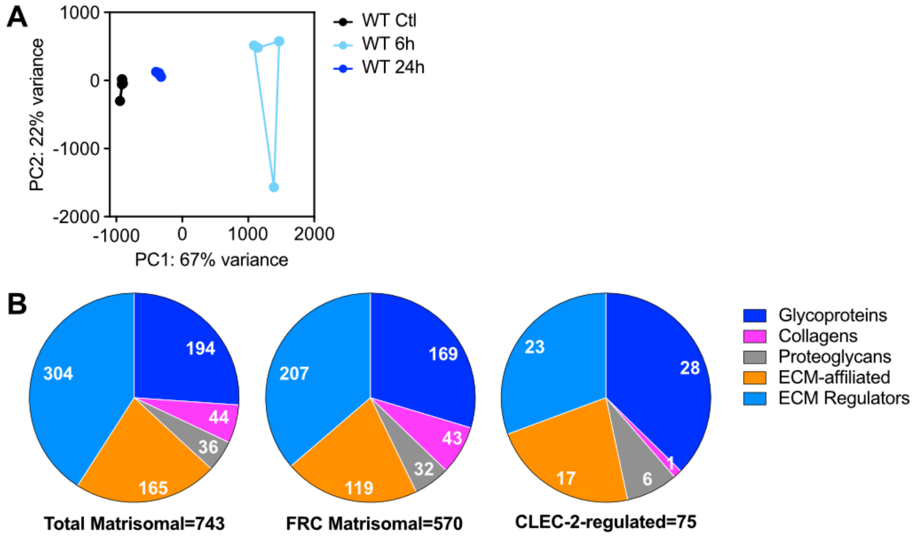


**Figure S1. Matrisomal genes regulated by CLEC-2.**

Gene expression by RNAseq in control FRCs treated with CLEC-2-Fc for 6 and 24 hours. A) CLEC-2-Fc-regulated gene (more or equal than 2-fold) cluster in a PCA space. B) Number of genes per category of matrisomal components in the indicated datasets.

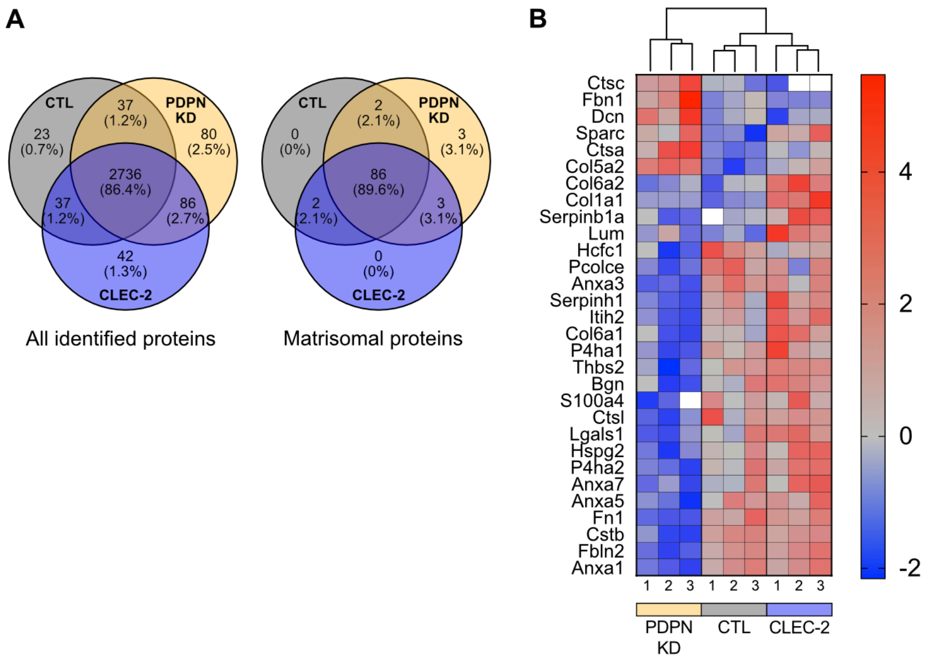


**Figure S2. Proteomics of FRC-derived matrices.**

*In vitro* FRC cell line-derived matrices generated after 5 days in culture were subjected to proteomic analysis by mass spectrometry. A) Venn diagrams showing overlap between cell lines of proteins detected. Areas shown are not proportional to percentage of overlap. C) Heatmap of matrisomal proteins significantly changed by one-way ANOVA, Turkey's multiple comparisons test. Three replicates for each condition automatically clustered are shown. Colour code represents z-scores. Not detected is represented by the white squares.

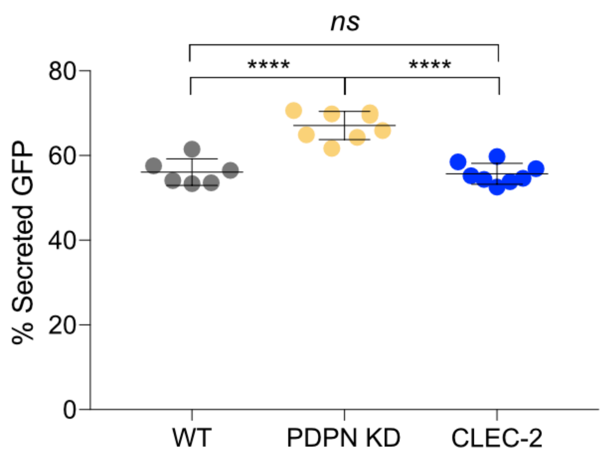


**Figure S3. Secretory activity in FRC cell lines.**

FRC cell lines were transfected to express GFP tagged for secretion. GFP levels in cell supernatants and lysates were determined by ELISA. Dot plot shows percentage of GFP in supernatant relative to total. Dots represent replicates form independent experiments (n=2). ****$P$<0.00005, one-way ANOVA, Turkey's multiple comparisons test. NS, not significant.

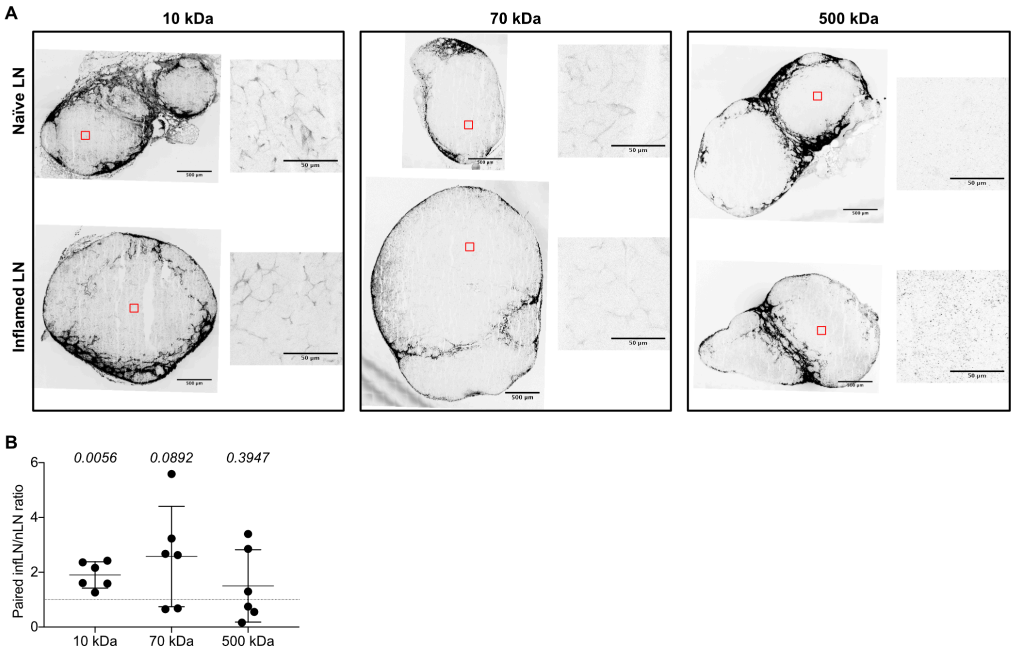


**Figure S4. Antigen uptake *in vivo*.**

Mice were immunized by subcutaneous injection of IFA/OVA on the right flank. 5 days later, fluorescently labelled dextrans with the indicated sizes were injected on both flanks. A) Immunofluorescence of 20 microns thick cryosections of naïve and inflamed draining LNs 30 minutes post dextran injection. Maximum z stack projections are shown of representative tile scans and zoomed areas are shown. B) Number of dextran-positive cells per draining LNs 90 minutes after dextran injection was determined by flow cytometry. Dot plot shows ratios between paired inflamed and naïve LNs from same individuals (n=6). *P*-values are shown for each dextran, unpaired t test.

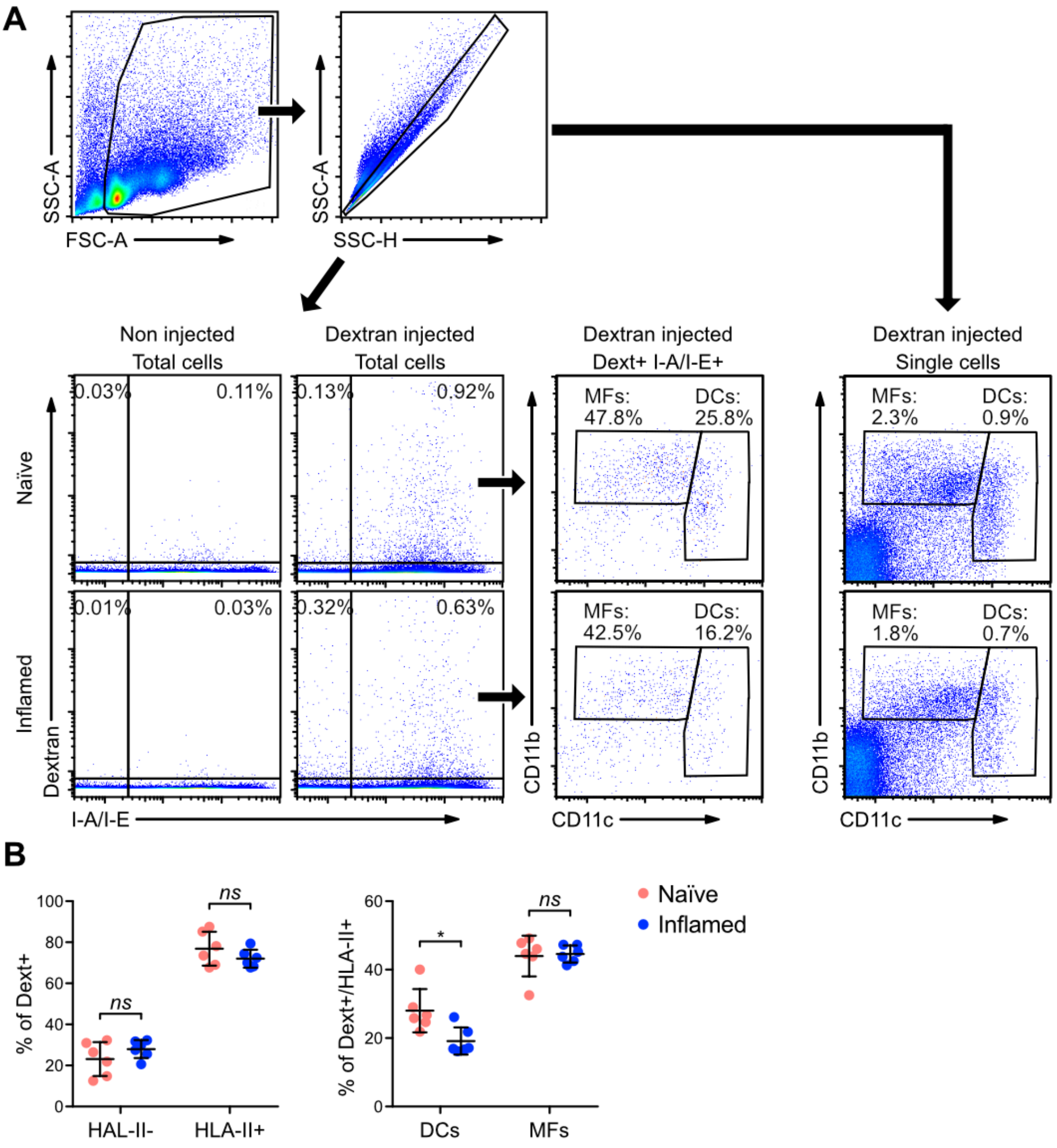


**Figure S5. Gating strategy and population analysis for in vivo antigen uptake.**

A) Representative dot plots showing gating strategy. Live cells were gated according to FSC/SSC parameters and doublets were excluded prior to analysis. Total cells were used to gate dendritic cell (DCs) and macrophage (MF) populations. B) Percentage of the indicated cell populations within the dextran-positive subset. Each dot represents an individual (n=6). *P<0.05, one-way ANOVA, Turkey's multiple comparisons test.

Table 1. Primary antibodies and imaging reagents used for immunofluorescence.

| Target | Isotype | Conjugated | Company | IF | Flow cytometry | WB | ELISA |
|---|---|---|---|---|---|---|---|
| α-Tubulin | Mouse monoclonal (236-10501) | Unconjugated | Thermo Fisher Scientific (A11126) | 1 µg/ml | | | |
| CD11b | Rat (M1/70) | APC/Cy7 | BioLegend (101225) | | 0.25 µg/106 cells | | |
| CD11c | Armenian Hamster (M1/70) | Brilliant Violet 605™ | BioLegend (117333) | | 0.25 µg/106 cells | | |
| Collagen I | Rabbit polyclonal | Unconjugated | Abcam (ab34710) | 10 µg/ml | | | |
| Collagen IV | Rabbit polyclonal | Unconjugated | Abcam (ab6588) | 10 µg/ml | | | |
| Collagen VI | Rabbit polyclonal | Unconjugated | Abcam (ab6588) | 10 µg/ml | | | |
| Fibronectin | Rabbit polyclonal | Unconjugated | Sigma-Aldrich (F3648) | 1.2 µg/ml | | | |
| F4/80 | Rat (BM8) | PerCP | BioLegend (123125) | | 1 µg/106 cells | | |
| GFP | Sheep polyclonal | Unconjugated | BioRad (4745-1051) | | | | (1ry) 0.1 µg/ml |
| GFP | Rabbit polyclonal | Unconjugated | Fisher Scientific (A-6455) | | | | (2ry) 1: 20000 |
| Histone H3 | Mouse monoclonal (mAbcam (24834) | Unconjugated | Abcam (ab24834) | | | 0.5 µg/ml | |
| Hoechst | | | Fisher Scientific (10150888) | 12.3 µg/ml | | | |
| I-A/I-E | Rat (M5/114.15.2) | Alexa Fluor® 647 | BioLegend (107617) | | 0.25 µg/106 cells | | |
| Phalloidin | | Alexa Fluor® 488 | Cell Signaling Technology (8878S) | 1x | | | |
| Phalloidin | | Alexa Fluor® 647 | Thermo Fisher Scientific (A22284) | 1x | | | |
| Phospho-Paxillin (Tyr118) | Rabbit polyclonal | Unconjugated | Cell Signaling Technology (2541S) | | | | |
| Podoplanin | Mouse monoclonal (811) | Unconjugated | Acris (DM3501) | 1 µg/ml | | | |
| Laminin | Rabbit polyclonal | Unconjugated | Abcam (ab11575) | 6.5 µg/ml | | | |
| LL5β | Mouse antisera | Unconjugated | kindly shared by Prof J. Sanes – Harvard University | 1:500 | | 1:500 | |
